# Cognitive modes involved in emotion regulation identified using Constrained Principal Component Analysis for fMRI (fMRI-CPCA)

**DOI:** 10.1101/2025.10.27.684758

**Authors:** Carien M. van Reekum, Emma Tupitsa, William K. Lloyd, Ava Momeni, John Shahki, Eva Feredoes, Todd S. Woodward

## Abstract

Meta-analyses of functional magnetic resonance imaging (fMRI) studies have identified networks of widely distributed brain regions supporting emotion regulation. These overlap with attentional or cognitive control brain networks. The literature is short on data speaking to specific neurocognitive functions of these broad brain networks in reappraisal - a key emotion regulatory strategy involving the reframing of an event according to a goal to increase or decrease experienced emotion. We address this gap by examining both the spatial configuration and temporal profile of event-related blood oxygenation level dependent (BOLD) responses during a task requiring reappraisal. We analysed fMRI datasets obtained from 84 participants (51% female) who were instructed to increase or decrease their emotional response to unpleasant images. We extracted spatial maps and their estimated temporal event-related BOLD signal changes of four components with the highest loadings. Neurocognitive functions were derived by mapping each component onto templates of previously identified task-based cognitive modes. This analysis yielded four cognitive modes: 1) “multiple demand” 2) “response”, 3) “re-evaluation”, and 4) “default mode". The temporal profiles showed particularly prominent patterns for the increase and decrease conditions in “multiple demand” (mode 1) and “re-evaluation” (mode 3) respectively. These findings highlight a central role for specific neurocognitive processes linked to attentional control (“multiple demand”) and switching (“re-evaluation”), as part of the broad brain networks supporting reappraisal. Moreover, the level of neural engagement of these cognitive modes varies depending on the regulatory goal. These findings provide tangible targets for neurocognitive interventions such as neurostimulation when emotion regulation is compromised.

## 1. Introduction

Emotion regulation supports mental health and wellbeing in everyday life. Dysregulated emotion is associated with various mental health conditions, including anxiety and depression (American Psychiatric Association, 2022). Given the relevance to our mental health, research into the neural mechanisms of emotion (dys)regulation expanded exponentially in the 2000s since the pioneering psychological research by Gross and colleagues (Gross, 1998). Of the emotion regulation processes that can be studied, reappraisal – a key element in cognitive behavioural therapy - has garnered the most attention particularly in the field of cognitive neuroscience. Reappraisal involves reframing an emotional situation or thought to change the intensity or quality of the emotion experienced in order to achieve a goal state.

Recent meta-analyses of functional magnetic resonance imaging (fMRI) studies (e.g. Buhle et al., 2014; Kohn et al., 2014; Morawetz et al., 2017) have identified networks of distributed brain regions supporting emotion regulation. Whilst the meta-analyses diverge in their analytic approaches, they converge in identifying the systematic engagement of, amongst others, frontal and parietal cortical brain areas in reappraisal across studies. In line with neurocognitive models of emotion regulation (Braunstein et al., 2017; Ochsner et al., 2012), these networks are widely thought to reflect top-down, cognitive control processes involved in the reframing of emotional stimuli, with corresponding changes in activity in brain areas, such as the anterior insula and amygdala, facilitating bodily responding^1^.

Two recent studies provide a step-change in our understanding of brain networks involved in emotion regulation. Using a meta-analytic clustering approach across fMRI datasets derived from a variety of emotion induction techniques and regulation strategies, Morawetz and colleagues (Morawetz et al., 2020) identified 4 distinct networks. Using an existing database (BrainMap, see https://brainmap.org/index.html), they deduced the functional correlates of these networks. This yielded two maps related to regulation, with associations with working memory, response inhibition, and language, and two maps more closely linked to emotion perception and generation, and bodily processes/awareness. Bo and colleagues (Bo et al., 2024) similarly provide evidence for overlapping and distinct networks involved in emotion generation and regulation. Based on consensus across two well-powered independent datasets, the authors identified several neural systems: one system that is shared between emotion generation (involving appraisal) and emotion regulation (involving reappraisal), and one specific to reappraisal-based regulation not shared with emotion generation processes. Matching the spatial component maps onto Neurosynth (Yarkoni et al., 2011), the study further highlights that the reappraisal-specific component map correlated closely with executive function (including response inhibition), as well as “empathy and interaction” topic maps (Bo et al., 2024, Fig. 6). This study underscores previous notions of the importance of cognitive control processes – and their corresponding networks – to reappraisal.

Despite these recent important advances in identifying functional brain networks supporting emotion regulation, our understanding of neurocognitive functions of the networks observed during a reappraisal task lacks in detail. Largely, the approaches taken thus far only consider spatial co-activation or convergence, without taking into account temporal information. Temporal information can particularly inform any differential engagement of a brain network as a function of different task conditions and cognitive processes during a task, and therefore aid the interpretation of network function. The field’s underspecification of network function is further hampered by the fact that a typical reappraisal task – where participants are asked to reframe an event depicted in order to change (intensify or dampen) their emotional experience - is comparatively unconstrained in terms of cognitive processes required when performing the task. Indeed, checks to ascertain that the task is performed as intended commonly rely on retrospectively asking participants to report on their strategies. Furthermore, the subtraction method generally employed in event-related fMRI implies that the control condition is closely matched to the experimental condition of interest. This is not always the case across a variety of tasks, and the reappraisal task is no exception. For the control condition, participants are typically asked to respond naturally, maintaining attention to the stimulus presented. This instruction is quite different from the request to reframe the stimulus presented in order to up- or downregulate emotion experienced, which requires an update of the initial evaluation. The lack of adequate experimental control leaves room for a variety of cognitive processes to be reflected in brain activation patterns observed. Indeed, past research has identified processes that at least partially explain reappraisal-linked fMRI findings, including attentional engagement/eye movements (van Reekum et al., 2007) and cognitive demand (De Voogd & Hermans, 2022; Urry et al., 2009). In summary, whilst the field has established various brain networks that are engaged in reappraisal, we know less about the specific functional contributions of these networks to reappraisal.

A potential solution is provided by Constrained Principal Component Analysis for fMRI data (fMRI-CPCA), developed by Woodward and colleagues (Metzak et al., 2011; Percival & Woodward, 2025). FMRI-CPCA is a data-driven method to identify task-based functionally connected networks and their associated temporal profile, by constraining the variance to the timing of the task events but without making any spatial or temporal assumptions on the evoked responses. The resulting components include both spatial maps (based on multivariate analyses) and estimates of the blood oxygenation level dependent (BOLD) signal changes (based on finite impulse response modelling) for each task condition. Each components’ combination of spatial maps and timing information can be subsequently ascribed to a “cognitive mode” ( Percival & Woodward, 2025). Cognitive modes can be defined as the cognitive (including sensory and motor) processes that reliably elicit BOLD signal pattern configurations across cognitive tasks and experimental conditions (Percival & Woodward, 2025). FMRI-CPCA has helped provide a finer specification of cognitive modes involved in, for instance, working memory (Sanford et al., 2020), lexical decision-making (Wong et al., 2020), and autobiographical cognition (Momeni et al., 2025), and has helped identify modes relevant to disease patterns in clinical samples, pertaining to hallucinations in schizophrenia (Roes et al., 2020) and disrupted motor network flexibility after stroke (Wadden et al., 2015).

The aim of the present study was to identify the cognitive modes involved in reappraisal by assessing task-evoked functional brain networks during emotion regulation. Participants engaged in a typical reappraisal task (Ochsner et al., 2004; Urry et al., 2006; see also Lloyd et al., 2021; Tupitsa et al., 2023 for details of the task employed here), where they were instructed to increase, maintain, or decrease their emotional response to negative images by reframing the situation depicted. Complementing recent work by others who identified brain networks associated with emotion generation vs regulation (Bo et al., 2024), we employed fMRI-CPCA to separate the spatial and temporal profile of cognitive modes involved in performing the task. Using template matching on prototypical examples of 12 cognitive modes (Percival et al., 2020), we then labelled the cognitive modes associated with the various neurocognitive processes engaged when performing reappraisal. This identification of task-general cognitive modes provides a data-driven, “objective discovery” approach to further our understanding of the role of neurocognitive processes in reappraisal, which can inform cognitive and neuroscience-based interventions to promote mental health and wellbeing.

## 2. Methods

### 2.1. Participants

91 adults (70 older adults, 21 younger adults) were recruited from the University of Reading’s Older Adult Research Panel via local newspaper and poster advertisements. All participants were right-handed and were screened for contraindication to MR scanning. No participants reported a history of neurological disorders or current use of steroid medication, and all participants scored between 25-30 on the Mini-Mental State Examination (MMSE), a range considered to reflect a “normal” cognitive function. All procedures were approved by the NHS Research Ethics Service and the University of Reading’s Research Ethics Committee. Participants provided written informed consent prior to their participation, and they received GBP7.50 per hour of financial compensation for their time. Due to leaving the study early (N=1), incomplete task data (N=3) and motion artifacts (N=3), the final sample size comprised 84 participants (64 older aged participants, M age = 69.7 years, range 54-84 years, 48% female, and 20 younger adults, M age = 27 years, range 18-34 years, 55% female).

### 2.2. Stimuli and tasks

#### 2.2.1. Stimuli

96 images were selected from the International Affective Picture System (Lang et al., 2008). Images were selected based on the normative valence ratings and categorised as either negative (76 images, valence ratings range = 1.46-3.85, M = 2.41, SD = 0.57) or neutral (24 images, valence ratings range = 4.38-5.88, M = 5.10, SD = 0.36). Picture categories were matched on luminosity, complexity and social content.

#### 2.2.2. Emotion regulation task

The voluntary emotion regulation task comprised the reinterpretation of negative events, such as scenes depicted in images or movie clips, to lessen or intensify an emotional response (cf. Urry et al., 2006; van Reekum et al., 2007). To induce negative affect prior to regulation, participants viewed each picture for a few seconds after which they were presented with an audio instruction to “suppress” (decrease), “enhance” (increase), or “maintain” (attend). Upon hearing “suppress” participants were asked to consider a less negative outcome of the scenario than they had thought prior to the instruction. For “enhance” they were asked to consider a more negative outcome, and for “maintain” they were asked to keep their original impression in mind. Participants received training and practice on the task immediately prior to scanning.

The full instructions provided during the training with respect to emotion regulation were: “On some trials we will ask you to either INCREASE ("enhance") or DECREASE ("suppress") the intensity of the emotion you are feeling in response to the picture. On other trials we will simply ask you to MAINTAIN ("maintain") the intensity of your emotion. When you hear the instruction “suppress”, we would like you to decrease the intensity of the emotion you are feeling when viewing the picture. You may do so by imagining a less negative outcome of the situation in the picture. As you are doing this, please continue to consider the situation to be real. On the other hand, when you hear the instruction “enhance”, we would like you to increase the intensity of the emotion you are feeling when you are viewing the picture. You may do this by imagining a worse outcome of the situation in the picture. Finally, when you hear the instruction “maintain”, please just maintain the intensity of the emotion you are feeling in response to the picture. You can do this by keeping in mind your initial reaction to the picture. Regardless of the instruction, you should keep focused on the content of the picture. In other words, please do not think of something unrelated to the picture, and do not try to generate an opposite emotion. For example, we ask you not to think of something comforting, like your home or a good friend, or think to yourself “it’s not real” to counteract any negative emotion you might experience. You may stop following the instruction ("enhance”, “suppress”, “maintain") as soon as the picture disappears from the screen.” Furthermore, example regulation strategies were suggested, e.g. for a picture of an injured animal to “enhance” their response they might consider the animal will die, or to “suppress” they might consider the animal will recover. Participants then practiced the task on 5 scenarios and were asked what strategy they used to follow the instruction. Training and practice were repeated if participants did not report the use of emotion regulatory strategies as intended.

Each picture was presented for 3 seconds before the audio instruction to regulate (“enhance”, “suppress” or “maintain”) was presented. The picture remained on the screen for a further 6 seconds, during which time participants were asked to keep following the instruction. Following the picture, a ratings screen was presented for 2 seconds, during which participants were asked to rate the prior image as neutral, somewhat negative, quite negative, or very negative by pressing a corresponding button on an MRI compatible button box held with their right hand. The 96 trials involved a pseudo-randomised presentation of picture-instruction combinations. All neutral pictures were accompanied by the “maintain” instruction.

The protocol was divided into four blocks, or “runs”, between which the participant may rest before commencing the next run. Inter trial interval (ITI) was jittered, and each block lasted for a duration of approximately 7 minutes. Details of the design are presented in Figure 1.

**Figure 1.**
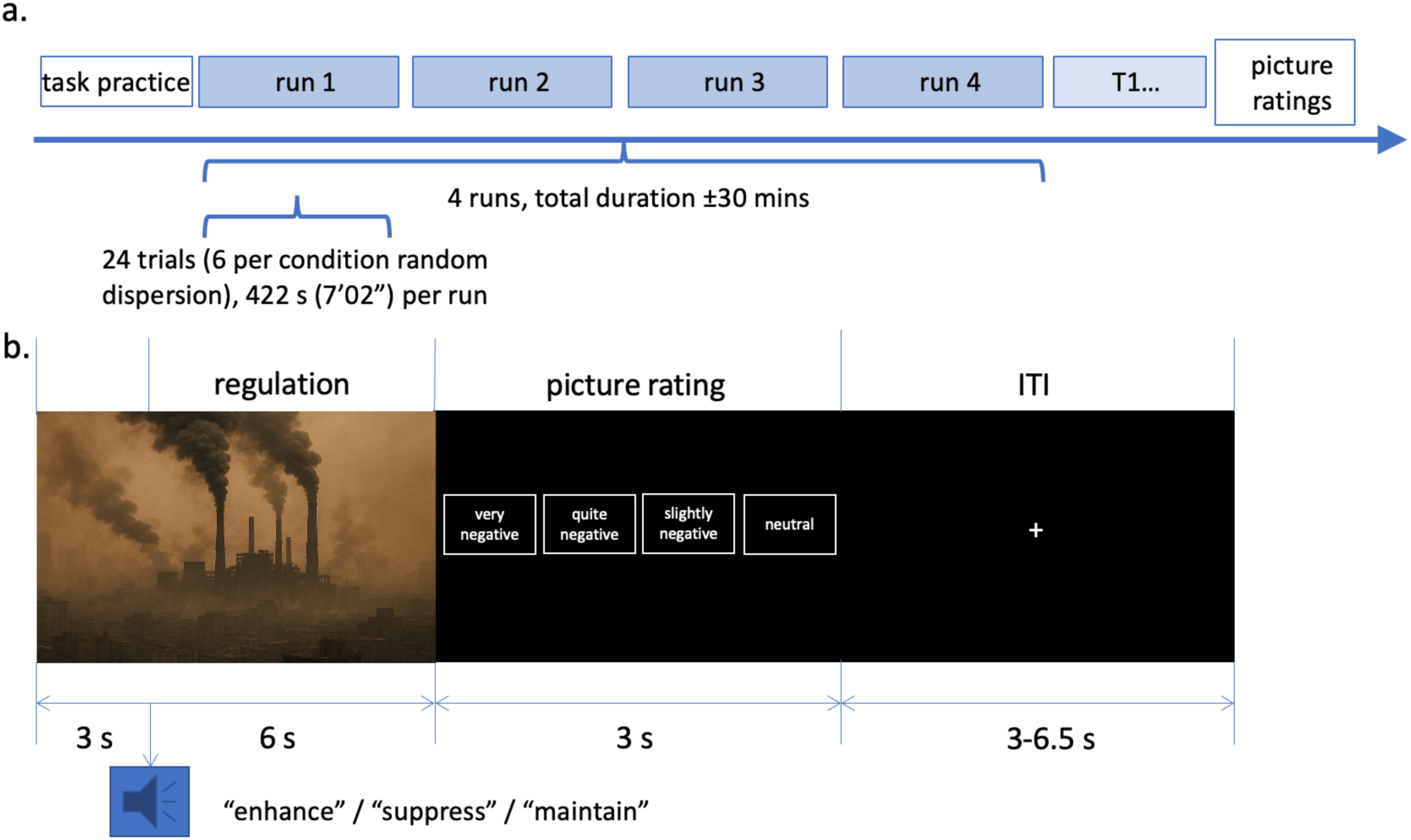
Task design. **a**. Order of study elements. Blocks in white (“task practice” and “picture ratings”) are elements that took place outside of the MRI unit. The in-scanner task comprised 4 runs, with 24 trials each. After the task was completed, a T1-weighted structural scan was obtained, alongside field maps, and a short task not reported here. **b**. Trial structure. The image was shown for 9 seconds in total, and the regulation instruction to increase (“enhance”), decrease (“suppress”) or to attend to the image without changing one’s feelings (“maintain”) was provided 3 seconds into the picture. Regulation co-terminated with the picture offset. A screen then prompted for a rating, presented for a fixed 3 s duration. For the intertrial interval (ITI), a screen displaying a fixation cross was then presented for a variable duration between 3-6.5 s. The image included here is generated by ChatGPT as a representative example of an image from the International Affective Picture System.

#### 2.2.3. Stimulus presentation

Stimuli were coded and presented using E-Prime (version 2.0.10.242, Psychology Software Tools, Inc., Pittsburgh, PA, USA) and delivered via VisualSystem binoculars (VisualSystem, NordicNeuroLab, Bergen, Norway). The VisualSystem binoculars were also used to monitor the participant’s eye movements and state of alertness throughout the experiment. Response to the ratings screen was recorded using a 4-button MR-compatible button box held in the participant’s right hand.

### 2.3 MRI data acquisition, reduction and analysis

#### 2.3.2. MRI procedure and image acquisition

Structural and BOLD functional imaging data were acquired using a 3T Siemens Magnetom scanner and 12-channel head coil (Siemens Healthcare, Erlangen, Germany) at the University of Reading’s Centre for Integrative Neuroscience and Neurodynamics. A 3D-structral MRI was acquired for each participant using a T1-weighted Magnetization Prepared Rapid Acquisition Gradient Echo (MPRAGE) sequence, with a repetition time (TR) = 2020 ms, echo time (TE) = 3.02 ms, inversion time (TI) = 900 ms, flip angle = 9°, field of view (FOV) = 250 x 250 x 192 mm, resolution = 1 mm isotropic, acceleration factor = 2, acquisition time = 9 min 7 s. Emotion regulation task-related fMRI data were collected in four identical blocks, using an echo planar imaging (EPI) sequence (211 whole-brain volumes, 30 sagittal slices, slice thickness = 3.0 mm, slice gap = 33%, TR = 2000 ms, TE = 30 ms, flip angle = 90°, FOV = 192 x 192 mm^2^, resolution = 3 mm isotropic, acquisition time = 7 min 2 s per block). A field map was acquired with a double-echo spoiled gradient echo sequence (TR = 488.0 ms, TE = 4.92/7.38 ms, voxel size: 3 × 3 × 3 mm^3^ (33% slice gap, flip angle 60°) that generated a magnitude image and 2 phase images. The field map image was computed from the 2 phase images. Further structural and functional MRI data were acquired as part of this protocol but are not reported here. The structural and emotion regulation fMRI task data are publicly available on OpenNeuro: https://openneuro.org/datasets/ds002620/versions/1.0.0

#### 2.3.3. fMRI preprocessing and registration

As described in Tupitsa et al. (2023), functional imaging data were pre-processed and analysed using FMRIB’s Software Library (FSL, version 6.0; www.fmrib.ox.ac.uk/fsl; (Jenkinson et al., 2012; Smith et al., 2004) and Analysis of Functional NeuroImages (AFNI, version 19.3.03; http://afni.nimh.nih.gov/afni; Cox, 1996). Initial pre-processing steps included skull stripping (non-brain removal) using FSL’s brain extraction tool (BET; Smith, 2002), motion correction using MCFLIRT (Jenkinson et al., 2002), field-map correction to correct for potential magnetic field inhomogeneity distortions, spatial smoothing using a Gaussian kernel with a full width at half maximum (FWHM) of 5mm, and high-pass temporal filtering (Gaussian-weighted least squares straight line fitting with sigma = 50 s). Each participant’s fMRI data was normalised to Montreal Neurological Institute (MNI) space via co-registration to their high resolution T1-weighted image.

Application of FSL’s MELODIC Independent Components Analysis (ICA; Beckmann & Smith, 2004) separated the fMRI BOLD signal into a set of spatial maps (independent components) representing neural signal and/or noise. Independent components containing structured temporal noise, including scanner and hardware artifacts, physiological artifacts (respiratory and/or cardiac noise), and motion-related noise were identified via visual inspection and removed using the FSL command line tool ‘fslregfilt’ for each emotion regulation task run (Griffanti et al., 2017). Following ICA filtering, low bandpass filtering was applied to the fMRI data using AFNI’s ‘3dBandpass’ tool to further remove confounding signals below 0.009 Hz and above 0.1 Hz. Prior to analysis, each participant’s corresponding mean functional timeseries image was added back to the bandpass filtered data using fslmaths to ensure compatibility with FSL’s FMRI Expert Analysis Tool (FEAT).

#### 2.3.4. fMRI-CPCA

Functional MRI data were analysed using fMRI-CPCA with varimax rotation (Metzak et al., 2011; Momeni et al., 2025; Woodward et al., 2006) (see the Appendix published in Momeni et al., 2025 for a comprehensive mathematical explanation of fMRI-CPCA). Generally, in fMRI-CPCA, first, multivariate multiple regression is used to constrain the variance in the BOLD data to that predictable from the task timings, specified by the finite impulse response (FIR) model. Then, dimension reduction via PCA is performed on the matrix of predicted scores. The PCA component loadings depict the dominant dimensions of intercorrelated voxel activity, and these can be classified into network prototypes (Percival, Zahid, & Woodward, 2020). The PCA component scores are regressed onto the FIR model matrix to obtain predictor weights that are used to: (1) produce plots depicting the BOLD changes for each component (i.e., each mode) in each condition at each time point for each participant, and (2) be submitted to a conventional statistical analysis (e.g. repeated-measures ANOVA). Thus, through fMRI-CPCA, one is able to (1) identify multiple BOLD networks that are simultaneously involved in the autobiographical event simulation task, (2) estimate the task-induced BOLD signal changes for each network at each time point in each condition for each participant, and (3) statistically test the effect of conditions on task-induced BOLD signal changes. The fMRI-CPCA application is openly available online (www.nitrc.org/projects/fmricpca).

The spatial maps of each component resulting from PCA are then matched with template maps. Whole-brain template images have been created for each of the fMRI task-based cognitive modes (Percival et al., 2020, Percival & Woodward, 2025) by averaging the images of several fMRI-CPCA components demonstrated to be excellent anatomical examples of a certain cognitive mode, in that they displayed distinctive and uniform anatomical and temporal activity patterns. An in-house template matching algorithm (Woodward et al., 2021) provides automatic classification information for any to-be-classified anatomical image to 12 task-based anatomical template images (11 distinct anatomical template images with separate classifications carried out for the one- and two-handed response modes). Each of these 12 template images has its own unique set of 20-30 brain slices that highlight highly replicable activity patterns (see Percival & Woodward, 2025). For each classification, the algorithm isolates these specific slices from the template and the to-be-matched image. The values in the template image and the component loadings in the to-be-classified image are correlated using Pearson’s *r* and then transformed to Fisher *Z* values. Peak activations in anatomical structures comprising these networks were localised using the Harvard-Oxford Cortical Structural Atlas (RRID:SCR_001476).

In this study, we estimated BOLD signal changes for 12 post-stimulus time points. Because scans were taken with a TR of 2s, this resulted in a model of the BOLD response over a 24-second window following the onset of the image and including the rating period and the ITI, whilst allowing for the hemodynamic delay (see Figure 1b). For this study, the data were modelled per run with 3 explanatory variables (EVs) used to describe the instruction conditions (decrease, increase, and maintain) with negative pictures, and 1 EV for the maintain instruction with neutral pictures. To determine the effects of regulation instruction type on network responses, we entered the subject-specific BOLD change curves estimated by fMRI-CPCA to a repeated-measures ANOVA (RM-ANOVA) using SPSS v.26. For each network extracted with fMRI-CPCA, we ran a 4 (Condition: Increase, Maintain, Decrease, Neutral) x 12 (time bins) repeated-measures ANOVA. Significant Condition x Time interaction effects are reported and followed up by a simple effects analysis (2×2 ANOVAs) and contrasts.

## 3. Results

### 3.1. Component selection

The scree plot indicated diminishing returns in variance explained beyond 4 components, hence spatial and temporal information of the first 4 components, explaining 18.8%, 11.6%, 4.3% and 2.8% respectively, were extracted for further analysis. As detailed above, the classification of each spatial map in cognitive modes was obtained by using an algorithm implemented in MATLAB to match the activity of each component onto a set of task-based network templates with distinct prototypes. The algorithm assigns Fisher’s Z-scores to each input brain network, reflecting how well the activity pattern matches to the template networks. A *Z*-score of 0.8 (corresponding to a correlation of 0.67) or above is considered “good”. The legends of Figures 2-5 provide the values of the highest Z-score obtained for each component map.

**Figure 2:**
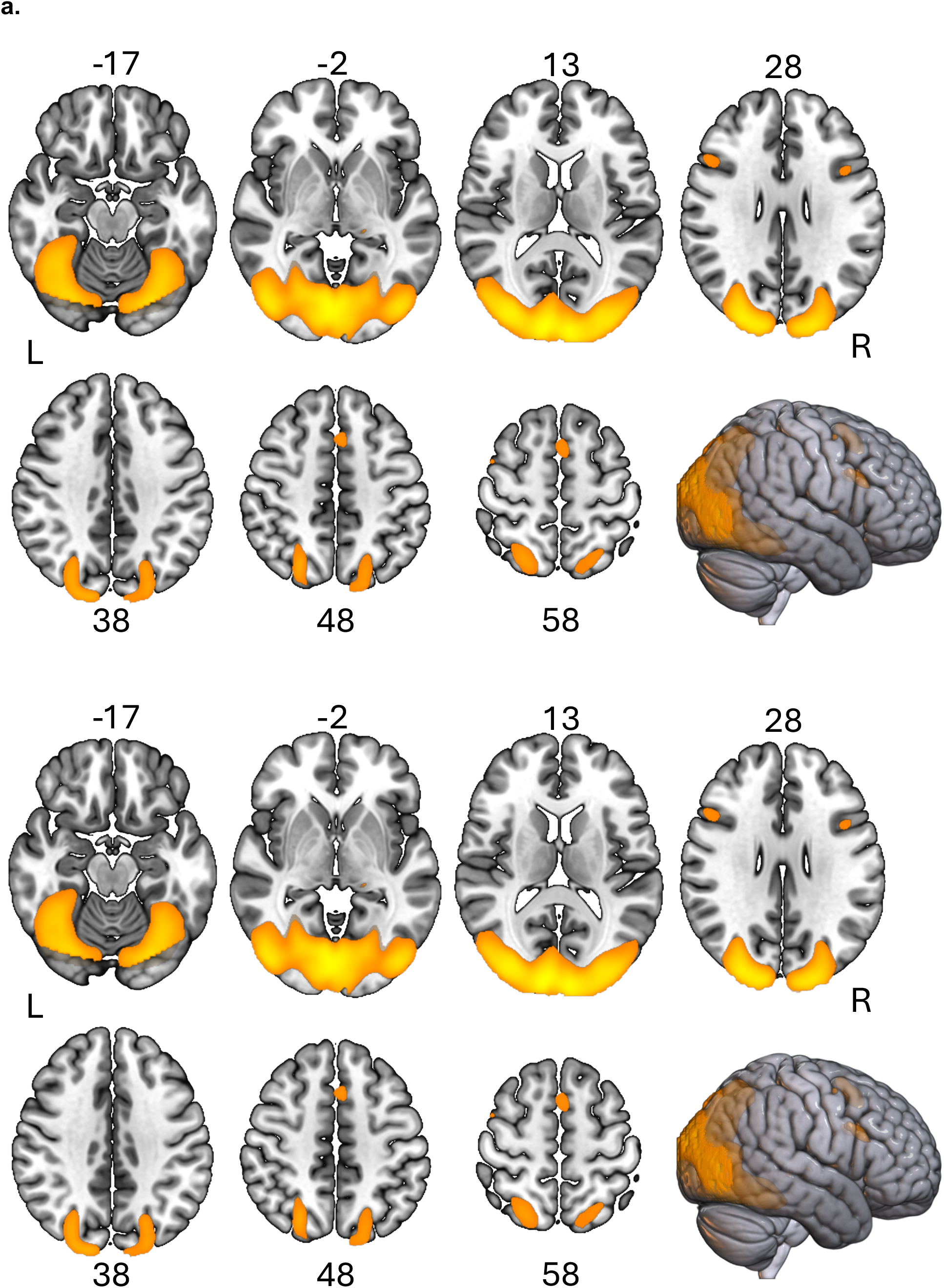
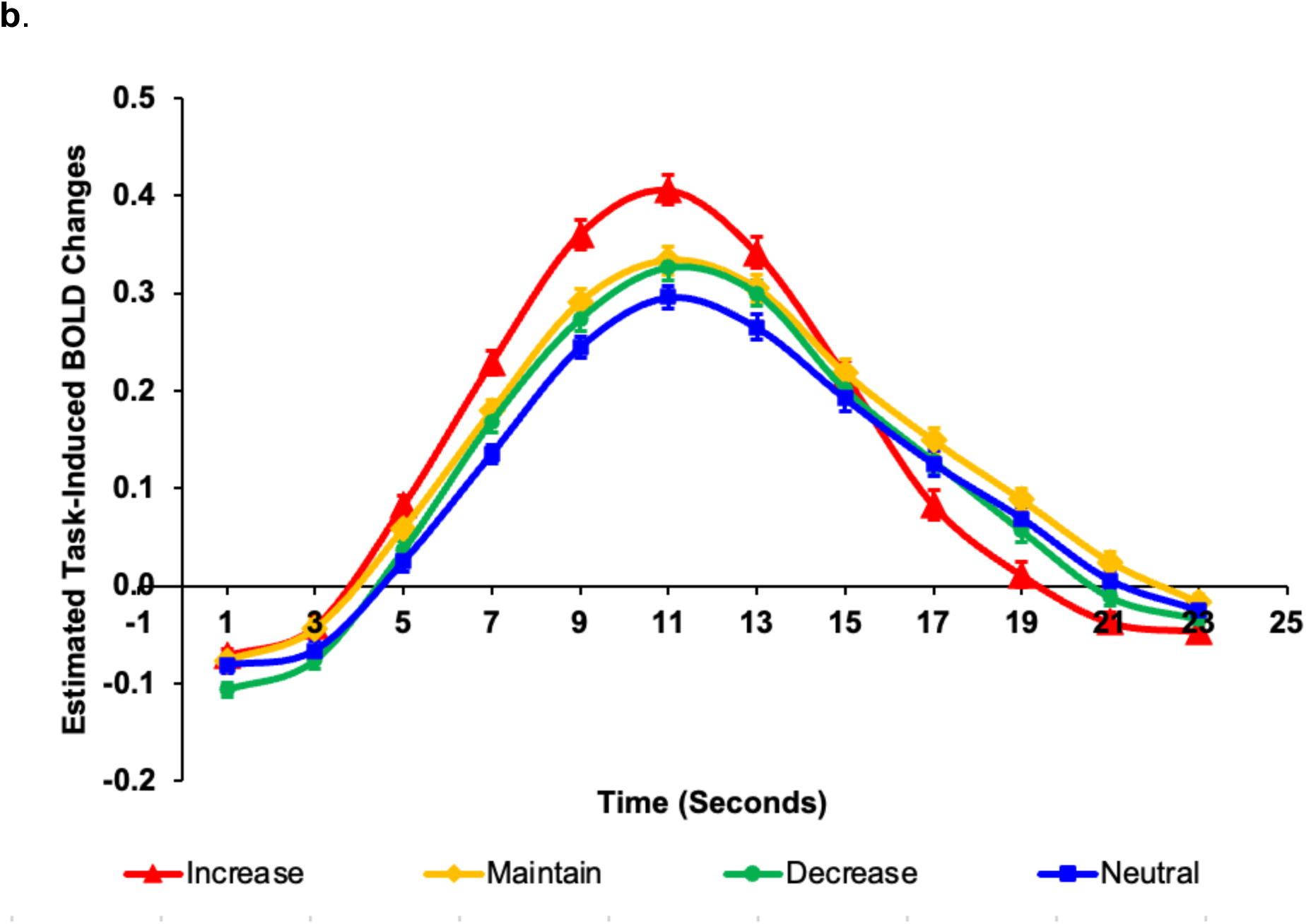
**a.** Dominant component loadings for Component 1, which mapped most strongly onto the multiple demand (MD) mode (Fisher’s Z=0.63) of the cognitive mode templates. The images are thresholded at the highest 10% of the component loadings (threshold = 0.24, maximum = 0.59). MNI Z-axis coordinates are displayed. L = left; R = right. **b**. BOLD signal change across the 12 2-second time bins, for each of the Increase (red), Decrease (green), and Maintain (yellow) regulation instruction conditions during negative images, and the Neutral (blue) image condition with the maintain instruction. Error bars depict standard errors.

Included in each Figure are the time courses of the finite impulse response-based predictor weights estimated for each 2-second (TR duration) time bin producing an estimated BOLD signal change for each of the 4 conditions. The estimated BOLD signal change time courses are estimated for each individual dataset, prior to merging across datasets. These time courses further help in determining which of the active regulation condition(s) engaged the functional network most strongly, relative to the negative and neutral “maintain” control conditions.

### 3.2. Components

#### 3.2.1. Component 1: Multiple Demand mode (MD)

As detailed in Table 1 (see end of document), clusters in this component involved bilateral activation in occipital pole (BA 17), fusiform gyrus (BA 18), lateral occipital cortex, (BA 19) extending into superior parietal cortex, supplementary motor area (SMA, BA 32), and inferior frontal gyrus, pars opercularis (BA 44). The spatial map best matched with the “multiple demand” (MD) mode, although the fit was moderate (Fisher’s *Z* = 0.63). As highlighted in recent work (Evora & Woodward, 2025; Wang et al., 2024), the MD mode and “initiation” (INIT) mode spatially overlap, but are distinguished by the peak of SMA activation in MNI’s axial slice 64. For MD, and as observed for the SMA cluster in this study, the peak is commonly lower, in slice 52 in the present study. The MD mode is derived from spatial maps obtained from a diverse range of tasks that all involved internal (re)construction, mentalising, information integration or reasoning following visual cues.

**Table 1.**
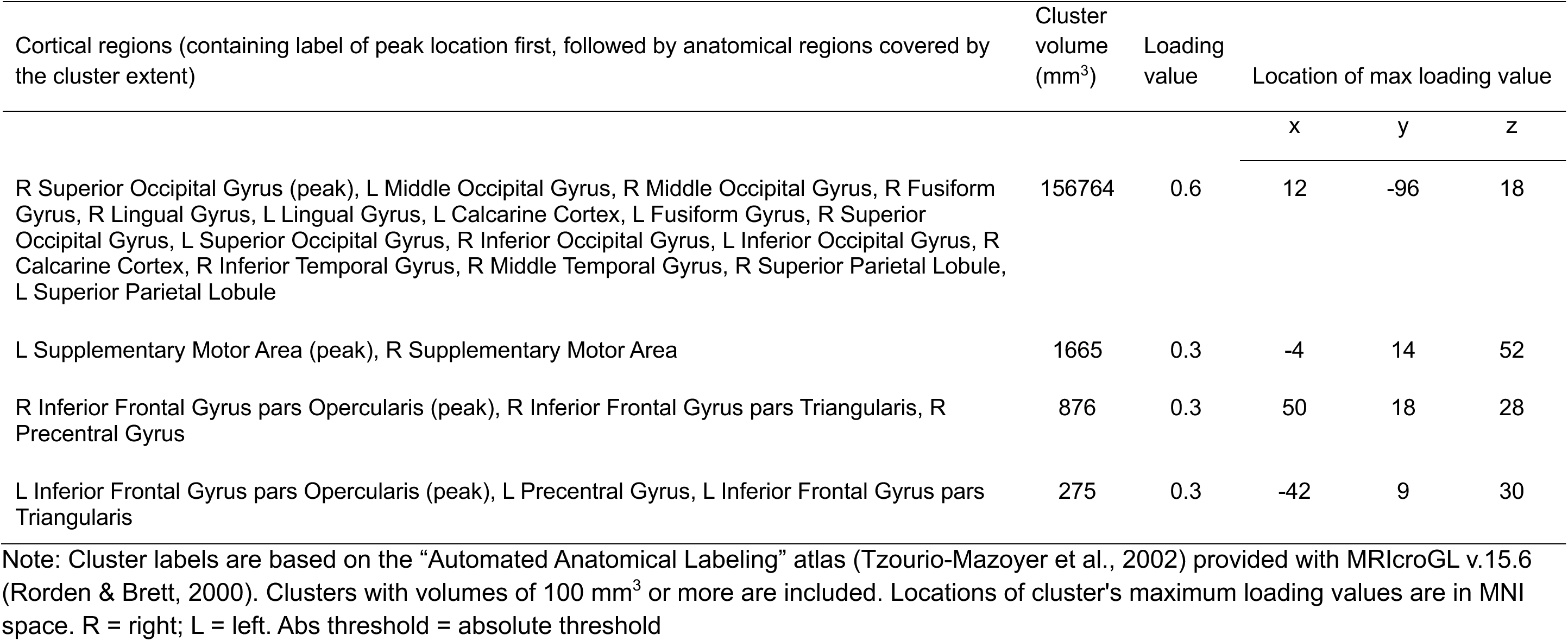
Cluster information for the top 10% loadings (abs threshold = 0.24) on Component 1 (Multiple Demand)

The 4 (Condition: Increase, Maintain, Decrease, Neutral) × 12 (Time) RM-ANOVA on the time courses derived from the HRF for each condition revealed a significant Condition × Time interaction, *F*(33,2739) = 20.62, *p* < .001, *η2p* = .20. Maintain and Decrease had very similar BOLD change patterns, and when only these two conditions were included in the ANOVA, the Condition × Time interaction was not significant *F*(11,913) = 1.59, *p* = .18. Therefore, to simplify interpretation of the Condition × Time interaction, these conditions were averaged together. However, the main effect of this analysis showed greater activity averaged over time for the Maintain condition (*M* = .13) relative to the Decrease condition (*M* = .10).

Ordering the conditions according to magnitude of the BOLD change (i.e., Increase > Maintain/Decrease combined > Neutral), and breaking down the Condition × Time interaction (p < .001) into 2 × 2 contrasts of adjacent time bins/conditions (with the latter ordered according to magnitude), the contrast of Increase > Maintain/Decrease combined was dominated by the increases from 3-9 s, and decreases from 11-17 s, and 21-23 s (all *p*s < .005), attributable to the higher peak and more rapid return to baseline for the Increase condition relative to Maintain/Decrease combined. The contrast of Maintain/Decrease > Neutral was dominated by the increases from 1-7 s and decreases from 13-15 s (all *p*s < .005), attributable to the Maintain/Decrease averaged condition having a higher peak than Neutral.

#### 3.2.2. Component 2: Response mode (RESP)

Anatomically, component 2 covered motor areas including bilateral – but with stronger activation in the left hemisphere - pre-motor and primary motor cortex, supplementary motor area (SMA) and pre-SMA, precuneus, insular cortex and cerebellar cortical activity (Fig. 3a, see Table 2 at the end of the document for full details). The spatial distribution is typical of motor responses such as button pressing in cognitive tasks, and indeed the map matched best with the Response (RESP) mode template, Fischer *Z* = 0.88 (Mascarenhas & Woodward, 2025). The RESP mode is further supported by the timing of the BOLD responses for this component, with a delayed peak at 17 seconds after image onset: Participants were prompted to provide a rating after picture offset,9 seconds into the trial. The fastest and highest activation at 17 seconds was observed for the neutral image condition (neutral-maintain), whilst the slowest (and more variable) response time was observed for the negative-increase task condition.

**Figure 3:**
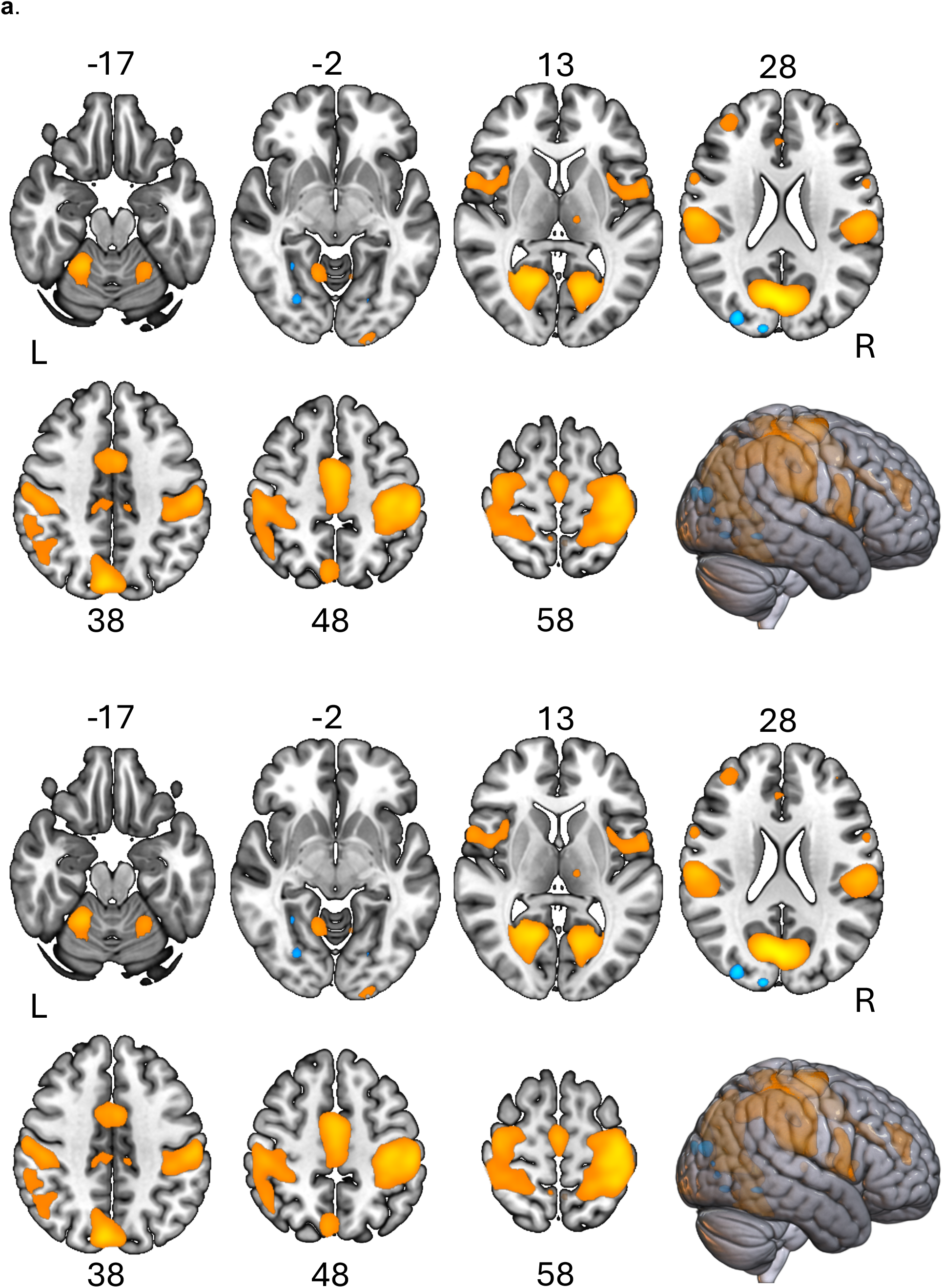
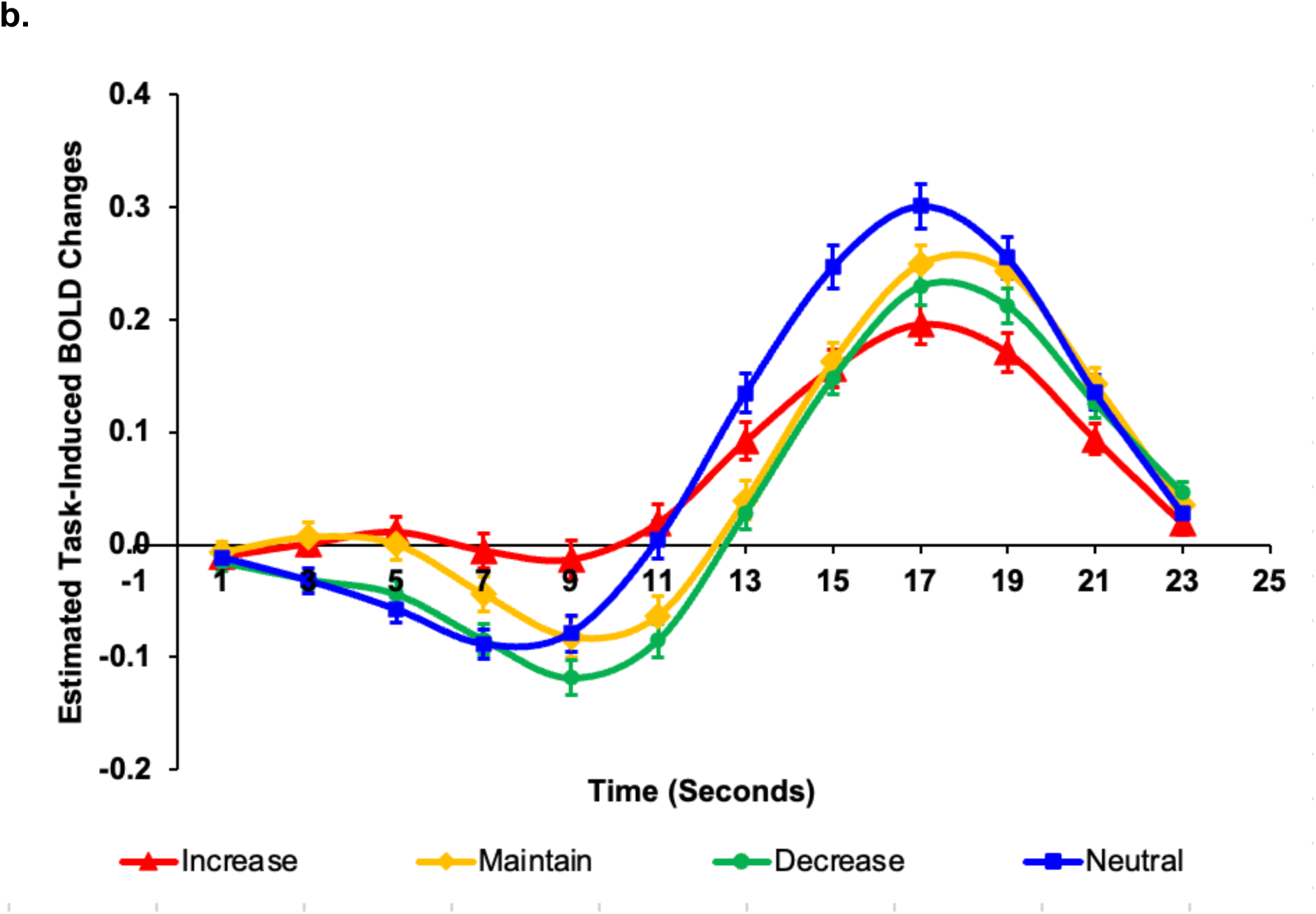
**a**. Dominant component loadings for Component 2, which mapped strongest on the response mode network (RESP) (*Z*=0.88) of all cognitive mode templates. The images are thresholded at the highest 10% of the component loadings (absolute threshold = 0.19, maximum = 0.42, minimum = −0.22). Negative voxels are depicted in blue. MNI Z-axis coordinates are displayed. L = left; R = right. **b**. BOLD signal change across the 12 2-second time bins, for each of the Increase (red), Decrease (green), and Maintain (yellow) regulation instruction conditions during negative images, and the Neutral (blue) image condition with the maintain instruction. Error bars depict standard errors.

**Table 2.** Cluster information for the top 10% loadings (abs threshold = 0.19) on Component 2 (Response)

| Cortical regions (containing label of peak location first, followed by anatomical regions covered by the cluster extent) | Cluster volume (mm<sup>3</sup>) | Loading value | Location of max loading value |  |  |
| --- | --- | --- | --- | --- | --- |
|  |  |  | x | y | z |
| L Cuneus (peak), R Calcarine Cortex, L Calcarine Cortex, R Cuneus, R Precuneus, R Lingual Gyrus, L Lingual Gyrus, R Cerebellum Lobule IV/V, R Cerebellum Lobule VI, L Precuneus, L Occipital Superior Gyrus, Midline Vermis Lobule VI | 43106 | 0.4 | 0 | -76 | 24 |
| L Postcentral Gyrus (peak), L Precentral Gyrus, L Inferior Parietal Lobule, L Supramarginal Gyrus, L Superior Parietal Lobule, L Precuneus, L Superior Temporal Gyrus, L Rolandic Opercular Area, L Superior Frontal Gyrus | 42673 | 0.4 | -40 | -22 | 64 |
| R Postcentral Gyrus (peak), R Supramarginal Gyrus, R Precentral Gyrus, R Inferior Parietal Lobule, R Superior Parietal Lobule, R Rolandic Opercular Area, R Superior Frontal Gyrus, R Angular Gyrus, R Superior Temporal Gyrus, R Precuneus | 36354 | 0.3 | 59 | -19 | 22 |
| R Middle Cingulate Cortex (peak), R Supplementary Motor Area, L Middle Cingulate Cortex, L Supplementary Motor Area, L Paracentral Lobule, R Anterior Cingulate Cortex, L Anterior Cingulate Cortex | 15763 | 0.3 | 0 | -6 | 54 |
| L Rolandic Opercular Area (peak), L Insular Cortex, L Precentral Gyrus, L Superior Temporal Gyrus, L Inferior Frontal Opercular Area, L Superior Temporal Pole, L Postcentral Gyrus | 4722 | 0.2 | -58 | 2 | 8 |
| R Rolandic Opercular Area (peak), R Insular Cortex, R Precentral Gyrus, R Inferior Frontal Opercular Area, R Superior Temporal Pole | 4634 | 0.2 | 59 | 7 | 8 |
| R Middle Frontal Gyrus | 2595 | 0.2 | 38 | 46 | 26 |
| L Cerebellum Lobule VI (peak), L Cerebellum Lobule IV/V | 775 | 0.2 | -23 | -54 | -22 |
| L Inferior Occipital Gyrus (peak), L Middle Occipital Gyrus, L Calcarine Cortex | 429 | 0.2 | -21 | -102 | -8 |
| L Middle Frontal Gyrus | 158 | 0.2 | -36 | 44 | 28 |
| L Thalamus | 139 | 0.2 | -12 | -20 | 8 |
| R Middle Occipital Gyrus (peak), R Superior Occipital Gyrus | 611 | -0.2 | 30 | -86 | 20 |
| R Cuneus (peak), R Superior Occipital Gyrus | 218 | -0.2 | 12 | -94 | 22 |
| R Fusiform Gyrus (peak), R Inferior Occipital Gyrus | 157 | -0.2 | 28 | -74 | -8 |
| L Lingual Gyrus (peak), R Lingual Gyrus, R Calcarine Cortex | 121 | -0.2 | 4 | -80 | 2 |

Note: Cluster labels are based on the “Automated Anatomical Labeling” atlas (Tzourio-Mazoyer et al., 2002) provided with MRICroGL v.15.6 (Rorden & Brett, 2000). Clusters with volumes of 100 mm<sup>3</sup> or more are included. Locations of cluster's maximum loading values are in MNI space. R = right; L = left. Abs threshold = absolute threshold

The 4 (Condition) × 12 (Time) RM-ANOVA revealed a significant Condition × Time interaction, *F*(33,2739) = 18.08, *p* < .001, *η2p* = .18. As with the previous component, Maintain and Decrease had very similar BOLD change patterns, but when only these two conditions were included in the ANOVA, the Condition × Time interaction remained significant *F*(11,913) = 2.58, *p* < .05, dominated by the decrease from 1-3 s for Decrease which is not present for Maintain. However, given the similarity of the response patterns and to simplify interpretation of the Condition × Time interaction, these conditions were averaged together.

Ordering the conditions according to magnitude of the BOLD change peak (i.e. Neutral > Maintain/Decrease combined > Increase), and breaking down the Condition × Time interaction (*p* < .001) into 2 × 2 contrasts of adjacent time bins/conditions (with the latter ordered according to magnitude), the contrast of Neutral > Maintain/Decrease was dominated by BOLD increases from 7-11 s, (all *p*s < .001), attributable to the earlier and higher peak for the Neutral condition relative to Maintain/Decrease. The contrast of Maintain/Decrease > Increase was dominated by the BOLD increases from 11-15s (all *p*s < .001), attributable to the Increase condition remaining flat and having a lower peak than Maintain/Decrease.

#### 3.2.3. Component 3: Re-Evaluation mode (RE-EV)

Activation clusters in this component (Fig. 4a, Table 3 at the end of the document) were observed in left and right lateral PFC extending from dorsal into orbitofrontal or frontal pole regions, medial PFC/SMA, and posterior cingulate cortex, bilateral superior/mid-frontal gyrus, inferior parietal cortex, and posterior mid-temporal cortex. The prototypical pattern for the re-evaluation mode was derived from a number of task-based studies including a “bias against disconfirmatory evidence” task and a task switching Stroop task where blocks of colour naming and word naming were alternated (Redway, Arreaza, et al., 2025). Activations in bilateral inferior temporal gyrus and cerebellar cortex, also typical for the re-evaluation network, were not observed in our dataset (“X marks the spot”, see Redway et al., 2024) likely due to slice positioning at acquisition, for which coverage of superior cortical regions was prioritised.

**Figure 4:**
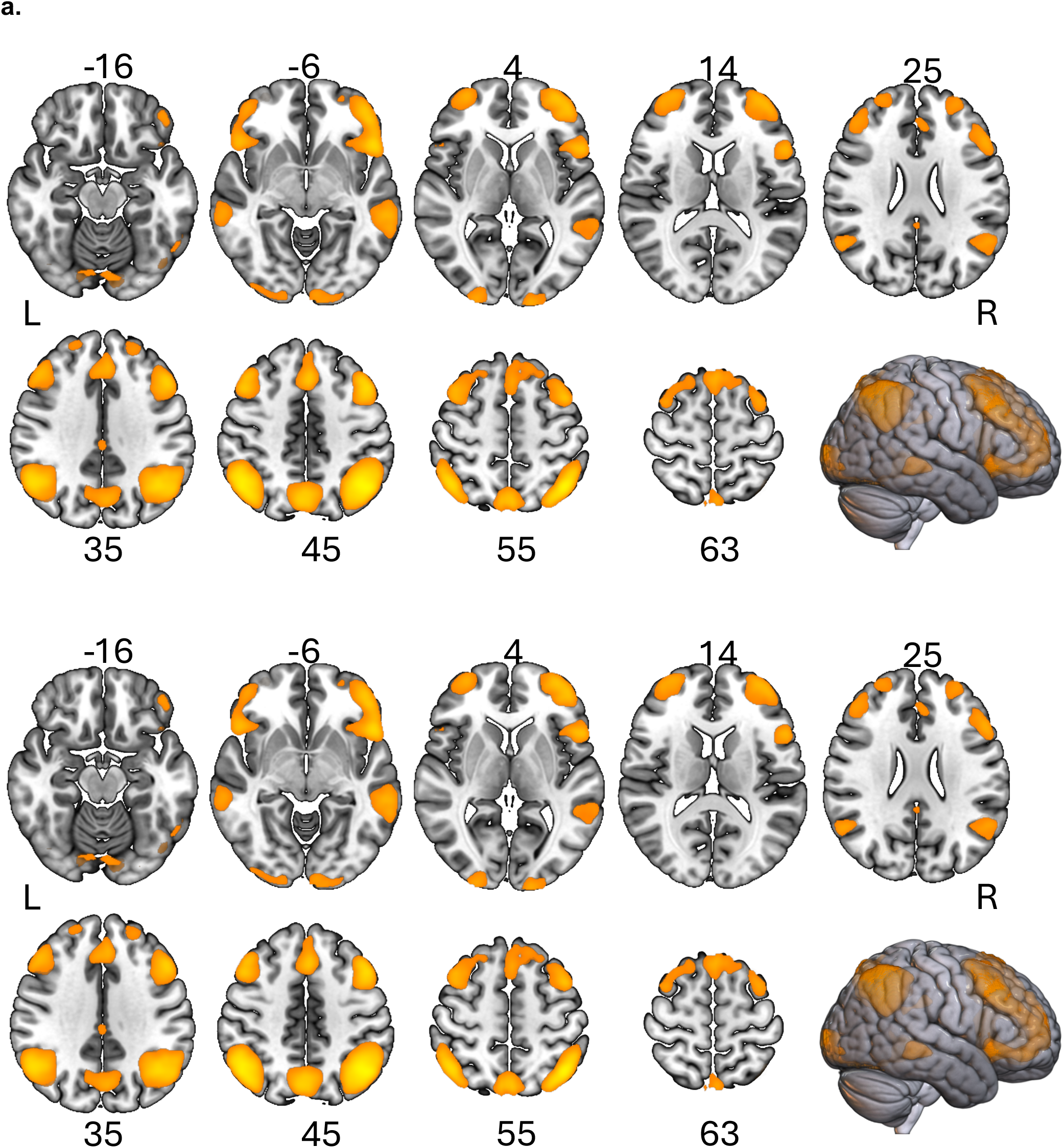
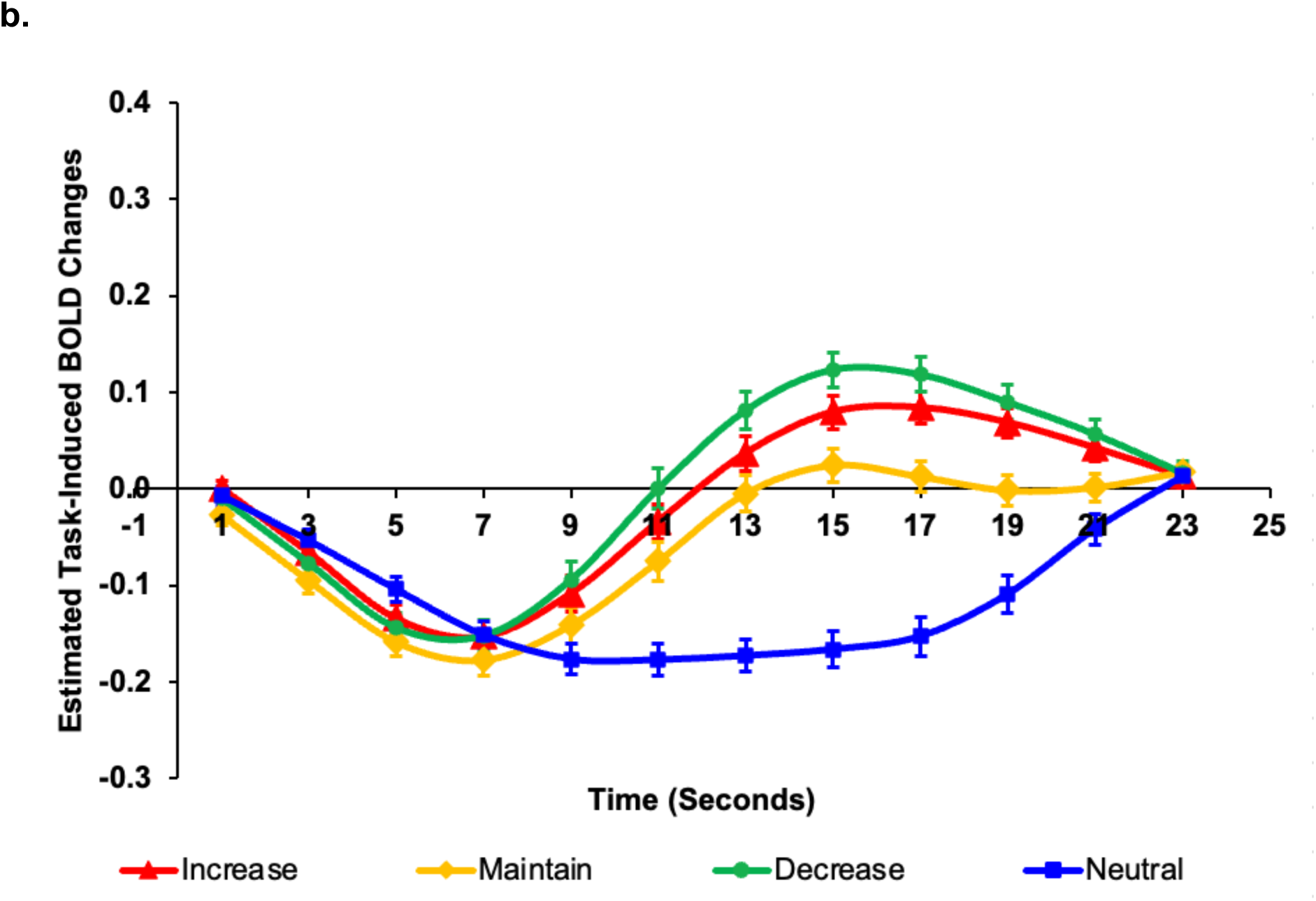
**a**. Dominant component loadings for Component 3, which mapped most strongly onto the re-evaluation network (RE-EV) (Z=1.16) of the “cognitive mode” templates in the independent database. The images are thresholded at the highest 10% of the component loadings (threshold = 0.12, maximum = 0.27). MNI Z-axis coordinates are displayed. **b**. BOLD signal change across the 12 2-second time bins, for each of the Increase (red), Decrease (green), and Maintain (yellow) regulation instruction conditions during negative images, and the Neutral (blue) image condition with the maintain instruction. Error bars depict standard errors.

**Table 3.** Cluster information for the top 10% loadings (abs threshold = 0.12) on Component 3 (re-evaluation)

| Cortical regions (containing label of peak location first, followed by anatomical regions covered by the cluster extent) | Cluster volume (mm<sup>3</sup>) | Loading value | Location of max loading value |  |  |
| --- | --- | --- | --- | --- | --- |
|  |  |  | x | y | z |
| L Inferior Parietal Lobule (peak), L Angular Gyrus, L Superior Parietal Lobule, L Middle Occipital Gyrus, L Supramarginal Gyrus, L Middle Temporal Gyrus | 26829 | 0.3 | -42 | -60 | 52 |
| R Angular Gyrus (peak), R Inferior Parietal Lobule, R Supramarginal Gyrus, R Superior Parietal Lobule, R Middle Occipital Gyrus, R Superior Occipital Gyrus, R Superior Temporal Gyrus | 20215 | 0.3 | 47 | -60 | 48 |
| L Middle Frontal Gyrus (peak), L Inferior Frontal Gyrus Triangular Part, L Medial Superior Frontal Gyrus, L Inferior Frontal Gyrus Orbital Part, L Supplementary Motor Area, L Superior Frontal Gyrus, L Inferior Frontal Gyrus Opercular Part, L Precentral Gyrus, L Orbital Middle Frontal Gyrus, L Insular Cortex, R Middle Cingulate Cortex, R Supplementary Motor Area | 59086 | 0.2 | -46 | 18 | 43 |
| R Middle Frontal Gyrus (peak), R Superior Frontal Gyrus, R Inferior Frontal Gyrus Orbital Part, R Inferior Frontal Gyrus pars Triangularis, R Orbital Middle Frontal Gyrus, R Inferior Frontal Gyrus pars Opercularis, R Insular Cortex, R Orbital Superior Frontal Gyrus, R Precentral Gyrus | 28969 | 0.2 | 46 | 26 | 40 |
| R Precuneus (peak), L Precuneus, L Cuneus, R Cuneus, R Superior Parietal Lobule | 13069 | 0.2 | 4 | -72 | 46 |
| L Middle Temporal Gyrus (peak), L Inferior Temporal Gyrus | 7060 | 0.2 | -58 | -36 | -4 |
| L Calcarine Cortex (peak), L Middle Occipital Gyrus, L Cerebellar Crus I, L Inferior Occipital Gyrus, R Cerebellar Lobule VI, L Lingual Gyrus, L Cerebellar Lobule VI, R Cerebellar Crus I, R Fusiform Gyrus | 4471 | 0.2 | -16 | -104 | 2 |
| R Calcarine Cortex (peak), R Inferior Occipital Gyrus, R Middle Occipital Gyrus, R Superior Occipital Gyrus, R Lingual Gyrus, R Cuneus | 2717 | 0.2 | 24 | -100 | 6 |
| R Middle Temporal Gyrus (peak), R Inferior Temporal Gyrus | 2204 | 0.2 | 62 | -33 | -4 |
| L Posterior Cingulate Cortex (peak), R Middle Cingulate Cortex, L Middle Cingulate Cortex, R Posterior Cingulate Cortex | 671 | 0.1 | -2 | -28 | 32 |
| R Cerebellar Lobule VI (peak), R Fusiform Gyrus, R Cerebellar Crus I | 155 | 0.1 | 36 | -68 | -20 |

Note: Cluster labels are based on the “Automated Anatomical Labeling” atlas (Tzourio-Mazoyer et al., 2002) provided with MRICroGL v.15.6 (Rorden & Brett, 2000). Clusters with volumes of 100 mm<sup>3</sup> or more are included. Locations of cluster's maximum loading values are in MNI space. R = right; L = left. Abs threshold = absolute threshold

The 4 (Condition) × 12 (Time) RM-ANOVA revealed a significant interaction of Condition × Time, *F*(33,2739) = 36.24, *p* < .001, *η2p* = .30. For this RE-EV mode, we compared each condition to the adjacent condition, starting at the highest -peaking condition (Decrease), and moving down to the lowest-peaking condition (Neutral). The highest peaking conditions, Decrease and Increase, produced a significant Condition × Time interaction when included together in the RM ANOVA, *F*(22,1826) = 2.95, *p* < .05, dominated by the increase from 9-11 s, which shows a steeper slope for Decrease relative to Increase, due to the higher eventual peak for Decrease relative to Increase. Comparing Increase to Maintain, this produced a significant Condition × Time interaction when included together in the RM ANOVA, *F*(11,913) = 3.00, *p* < .05, dominated by the change from 19-23 s, due to a flat slope for Maintain relative to Increase. This could be attributable to the return back to baseline require for Increase not present for Maintain, since the latter only recovered from deactivation (maximum at time point 7 s), but didn’t activate above baseline. Comparing Maintain to Neutral, this produced a significant Condition × Time interaction when included together in the RM ANOVA, *F*(11,913) = 45.94, *p* < .001, dominated by the change from 5-13 s and 17-23 s, due the fact that the Neutral did not recover from deactivation below baseline whereas Maintain did.

In all, attending to neutral images disengaged this mode; whereas decreasing one’s emotional response to negative pictures engaged this re-evaluation network the most, followed by increasing negative affect.

#### 3.2.4. Component 4: Default Mode B (DMB)

The final component includes clusters in regions typically associated with the “default mode network” (Raichle et al., 2001), i.e. orbitofrontal cortex, frontal pole extending into (ventro)medial prefrontal cortex, posterior cingulate cortex extending into precuneus, bilateral insula, and medial temporal cortex (see Fig 5, and Table 4 at the end of the document). The cognitive mode templates include two default mode networks, Default Mode A and Default Mode B, which show spatial similarity but are functionally distinct. For this study, the spatial pattern of component 4 matched best with Default Mode B (DMB), although Fisher’s *Z* suggests only a moderately good fit (*Z* = 0.50). Regions in this network characteristically show deactivation during psychological tasks. As discussed elsewhere (Ni et al., 2025; Redway, Jian, et al., 2025), some of these DMB regions are frequently included in other functional networks, notably somatosensory networks (e.g. insula), and visual and/or attentional networks (temporal and occipital cortex regions, fusiform gyrus), though showing activation rather than deactivation.

**Figure 5:**
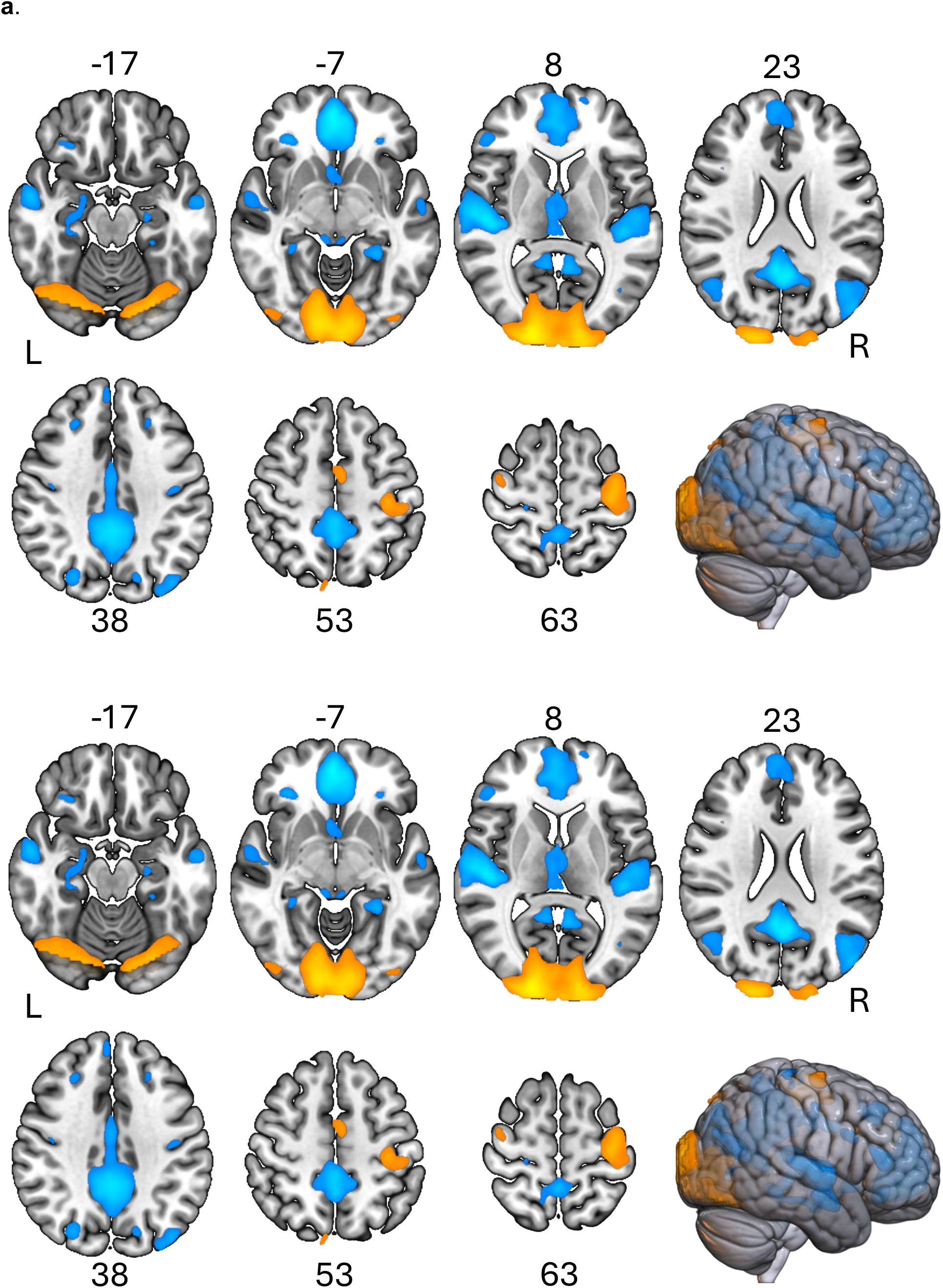
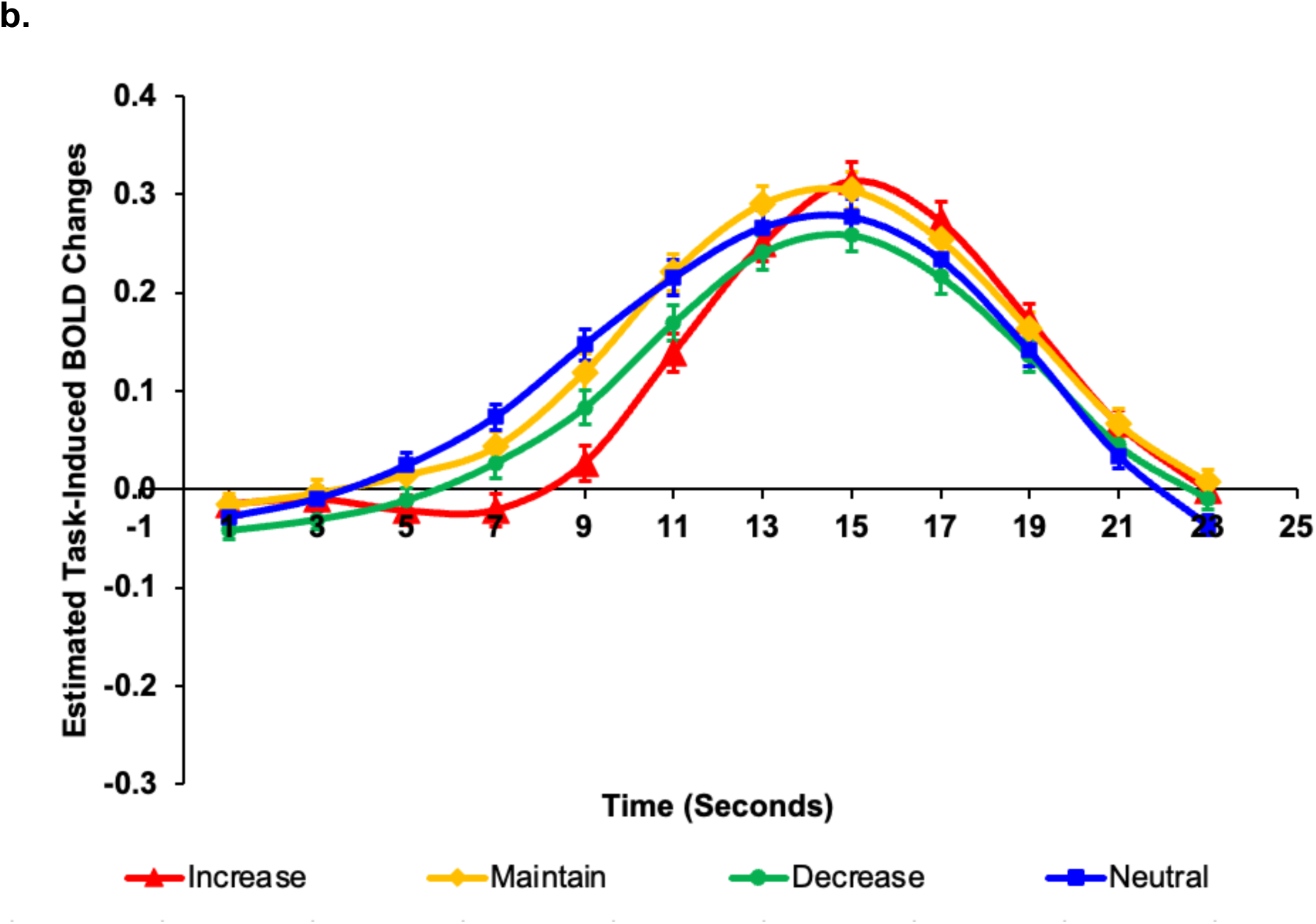
**a**. Dominant component loadings for Component 4, which mapped most strongly onto the default mode network type B (DMB) (*Z*=0.50) of the “cognitive mode” templates in the independent database. The images are thresholded at the highest 10% of the component loadings (absolute threshold = 0.09, maximum = 0.24, minimums = 0.19). Positive voxels are depicted in shades of yellow/orange, negative voxels are depicted in shades of blue. MNI Z-axis coordinates are displayed. **b**. BOLD signal change across the 12 2-second time bins, for each of the Increase (red), Decrease (green), and Maintain (yellow) regulation instruction conditions during negative images, and the Neutral (blue) image condition with the maintain instruction. Error bars depict standard errors.

**Table 4.** Cluster information for the top 10% loadings (abs threshold = 0.09) on Component 4 (Default Mode B)

| Cortical regions (containing label of peak location first, followed by anatomical regions covered by the cluster extent) | Cluster volume (mm<sup>3</sup>) | Loading value | Location of max loading value |  |  |
| --- | --- | --- | --- | --- | --- |
|  |  |  | x | y | z |
| R Precuneus (peak), L Precuneus, R Middle Cingulate Cortex, L Middle Cingulate Cortex, L Posterior Cingulate Cortex, L Calcarine Cortex, R Posterior Cingulate Cortex, R Calcarine Cortex, L Cuneus, R Paracentral Lobule, R Lingual Gyrus, | 35959 | -0.2 | 6 | -54 | 16 |
| R Medial Orbital Frontal Gyrus (peak), L Anterior Cingulate Cortex, R Superior Medial Frontal Gyrus, L Superior Medial Frontal Gyrus, R Anterior Cingulate Cortex, L Medial Orbital Frontal Gyrus, -(4) L Gyrus Rectus, R Gyrus Rectus, L Superior Frontal Gyrus, | 26036 | -0.2 | 2 | 52 | -6 |
| R Superior Temporal Gyrus (peak), R Middle Temporal Gyrus, R Heschl's Gyrus, R Rolandic Operculum, R Insular Cortex | 13825 | -0.2 | 50 | -18 | 6 |
| L Superior Temporal Gyrus (peak), L Middle Temporal Gyrus, L Heschl's Gyrus, L Rolandic Operculum, | 8508 | -0.2 | -40 | -33 | 12 |
| R Inferior Orbital Frontal Gyrus (peak), R Middle Orbital Frontal Gyrus, R Superior Orbital Frontal Gyrus, | 1390 | -0.2 | 32 | 34 | -12 |
| L Middle Occipital Gyrus (peak), L Middle Temporal Gyrus, L Angular Gyrus, L Inferior Parietal Lobule, | 7297 | -0.1 | -48 | -80 | 23 |
| R Thalamus (peak), L Thalamus, Vermis, R Lingual Gyrus, | 3434 | -0.1 | 6 | -32 | -4 |
| R Middle Temporal Gyrus (peak), R Middle Occipital Gyrus, R Angular Gyrus, | 2807 | -0.1 | 48 | -64 | 20 |
| R Parahippocampal Gyrus (peak), R Hippocampus, R Fusiform Gyrus, R Amygdala, | 1846 | -0.1 | 32 | -39 | -10 |
| R Superior Occipital Gyrus (peak), R Middle Occipital Gyrus, R Cuneus, R Superior Parietal Lobule, | 1526 | -0.1 | 26 | -76 | 42 |
| R Inferior Frontal Gyrus pars Triangularis | 1526 | -0.1 | 48 | 35 | 14 |
| R Middle Frontal Gyrus, R Superior Frontal Gyrus, | 1406 | -0.1 | 26 | 30 | 44 |
| L Fusiform Gyrus (peak), L Parahippocampal Gyrus, L Lingual Gyrus, | 1298 | -0.1 | -28 | -41 | -10 |
| L Middle Frontal Gyrus (peak), L Superior Frontal Gyrus, | 843 | -0.1 | -26 | 28 | 44 |
| R Postcentral Gyrus (peak), R Precentral Gyrus | 713 | -0.1 | 34 | -30 | 72 |
| L Inferior Orbital Frontal Gyrus (peak), L Middle Orbital Frontal Gyrus, | 665 | -0.1 | -30 | 34 | -11 |
| R Inferior Frontal Gyrus pars Opercularis (peak), R Precentral Gyrus, R Inferior Frontal Gyrus pars Triangularis, | 593 | -0.1 | 42 | 12 | 29 |
| R Caudate Nucleus (peak), L Caudate Nucleus, R Olfactory Cortex, L Olfactory Cortex, | 565 | -0.1 | 4 | 12 | -6 |
| R Precentral Gyrus (peak), R Postcentral Gyrus | 454 | -0.1 | 50 | -12 | 50 |
| L Cuneus (peak), L Superior Occipital Gyrus, | 322 | -0.1 | -16 | -76 | 40 |
| L Postcentral Gyrus | 292 | -0.1 | -43 | -16 | 36 |
| L Middle Occipital Gyrus | 197 | -0.1 | -42 | -68 | 4 |
| L Hippocampus | 158 | -0.1 | -24 | -18 | -17 |
| R Paracentral Lobule (peak), R Supplementary Motor Area, L Paracentral Lobule, | 105 | -0.1 | 2 | -25 | 74 |
| L Supplementary Motor Area (peak), R Supplementary Motor Area, L Middle Cingulate Cortex, | 605 | 0.1 | -4 | -6 | 54 |
| R Precentral Gyrus (peak), R Superior Frontal Gyrus | 659 | 0.1 | 40 | -11 | 66 |
| L Precentral Gyrus (peak), L Postcentral Gyrus, L Superior Frontal Gyrus, L Inferior Parietal Lobule, | 7055 | 0.2 | -37 | -14 | 68 |
| R Cuneus (peak), L Calcarine Cortex, R Calcarine Cortex, R Lingual Gyrus, L Lingual Gyrus, L Middle Occipital Gyrus, R Inferior Occipital Gyrus, L Superior Occipital Gyrus, R Fusiform Gyrus, R Superior Occipital Gyrus, L Fusiform Gyrus, R Middle Occipital Gyrus, R Cerebellum Lobule 6, L Inferior Occipital Gyrus | 55504 | 0.2 | 18 | -98 | 8 |

Note: Cluster labels are based on the “Automated Anatomical Labeling” atlas (Tzourio-Mazoyer et al., 2002) provided with MRICroGL v.15.6 (Rorden & Brett, 2000). Clusters with volumes of 100 mm<sup>3</sup> or more are included. Locations of cluster's maximum loading values are in MNI space. R = right; L = left. Abs threshold = absolute threshold

The 4 (Condition) × 12 (Time) RM-ANOVA revealed a significant interaction, *F*(33,2739) = 7.98, *p* < .001, *η2p* = .09. Maintain and Decrease had very similar BOLD change patterns, and when only these two conditions were included in the ANOVA, the Condition × Time interaction was not significant *F*(11,913) = 1.21, *p* = .31, so to simplify interpretation of the Condition × Time interaction, these conditions were averaged together. However, the main effect of this analysis showed greater activity averaged over time for the Maintain condition (*M* = .12) relative to the Decrease condition (*M* = .09). Ordering the conditions according to magnitude of the BOLD change peak (i.e. Increase > Maintain/Decrease > Neutral), and breaking down the Condition × Time interaction (*p* < .001) into 2 × 2 contrasts of adjacent time bins/conditions (with the latter ordered according to magnitude), the contrast of Increase > Maintain/Decrease combined was dominated by differences during 3-7 s and 11-15 s (all *p*s < .001), attributable to the higher peak and later increase to peak for the Increase condition relative to Maintain/Decrease combined. The contrast of Maintain/Decrease > Neutral was dominated by the increases from 9-13 s (all *p*s < .05), attributable to the Neutral condition starting activation earlier than Maintain/Decrease.

## 4. Discussion

Emotion regulation is a core ingredient of emotional wellbeing, and cognitive neuroscience has focused on reappraisal as a key emotion regulation process. Recent meta-analyses largely converge on the identification of cortical neural systems supporting reappraisal, a complex process composed of diverse cognitive operations. Despite recent advances in our understanding of which brain systems are crucially – and uniquely – involved in reappraisal (e.g. Bo et al., 2024; Morawetz et al., 2020), the neurocognitive components of reappraisal remain loosely defined. Using fMRI-CPCA, a data-driven method to cluster spatiotemporal fMRI patterns, we aimed to decompose brain-wide activity over the time-course of reappraisal into patterns that could be associated with cognitive subcomponents underlying the process. .Participants performed a typical reappraisal task, and results provide evidence for the involvement of four distinct patterns subserving the following cognitive modes: 1) a “multiple demand” mode corresponding to brain maps derived from studies employing visual cues and requiring high-level attention and cognitive elaboration of various elements; 2) a “response” mode largely reflecting motor responses engaged during the provision of ratings; 3) most strikingly, a “re-evaluation” mode derived from brain imaging studies using non-affective tasks requiring evaluation (confirm or disconfirm) of information and to switch task goals; and finally 4) a “default mode” largely comprising a task-negative network.

The reappraisal task typically involves one or two explicitly instructed, active regulation conditions, where individuals increase or decrease their emotional response to emotion-eliciting material, often negative pictures. Early brain imaging research in emotion regulation suggested that regulatory goal was associated with somewhat different “cognitive control” patterns of cortical brain activation (e.g. Ochsner et al., 2004; Urry et al., 2009), since confirmed in an fMRI meta-analysis separating goals, strategies and emotion induction material (Morawetz et al., 2017). Two recent studies pushed the envelope in our understanding of the function of distinct frontal, frontoparieto-insular and frontotemporal brain networks in emotion regulation, separating emotion generation, and common appraisal, from reappraisal networks (Bo et al., 2024; Morawetz et al., 2020). Both studies also converged in using existing databases to label spatial maps obtained for reappraisal with cognitive functions, identifying response inhibition, cognitive conflict, and language. Yet, both studies aggregated across regulatory goals, leaving the neurocognitive functions that may support each regulatory goal – as suggested by earlier work (e.g. Ochsner et al., 2004; Urry et al., 2009) - unaddressed. The findings presented here fill this gap and indicate that increasing and decreasing negative affect strongly engage the “multiple demand” and “re-evaluation” cognitive modes. Crucially, the temporal profiles highlight that the goal to increase negative affect demonstrated the highest BOLD response for the multiple demand mode, whilst the goal to decrease negative affect showed the highest BOLD response in the re-evaluation mode. Thus, whilst the goal to increase or decrease negative affect both engaged “cognitive control” networks, these regulatory goals diverge in their strength of engagement of each mode. This finding underscores that different cognitive operations are at play when regulating emotion to achieve a desired emotional outcome, reflected in levels of engagement of different neural systems. We will next briefly discuss findings for each cognitive mode separately.

### Multiple Demand (MD)

As summarised by Wang and colleagues (Wang et al., 2025), the MD mode is one of several modes engaging brain regions associated with visual attention, and has been observed across a number of task domains requiring different types of cognitive demand. Prior work by Duncan and colleagues has identified a distinct network involving regions widespread across the cortex, but loading on the “frontoparietal network”, which they named the “multiple demand system” (Duncan, 2010). They propose that the multiple demand system supports attentional integration or the synthesis of components of a cognitive operation (Shashidhara et al., 2024). With fMRI-CPCA, Wang and colleagues (2024) highlight that MD is not the only mode observed in tasks involving visual stimuli. And while the peaks of the MD mode derived from fMRI-CPCA overlap with regions in the visual network, ventral attention network, and the frontoparietal network identified by others (e.g. Yeo et al., 2011), so do some of the other cognitive modes identified (Percival & Woodward, 2025).

In this study, MD predominantly captured posterior visual attention brain areas, and the temporal profiles indicated that the responses started to diverge around 7 seconds after image onset, likely due to the regulation instruction provided a few seconds into the picture presentation. The regulatory goal to increase negative affect was associated with the highest response modulation at the peak (at 11 s), while the decrease condition demonstrated similar magnitudes at the peak to the maintain condition. This finding suggests that the increase condition required more multiple demand or attentional integration than the decrease and attend condition. Indeed, past research using eye tracking has demonstrated that people engage in redirecting visual attentional processes on the emotionally evocative elements of the images when instructed to increase their experienced affect, relative to the attend control condition and relative to when the goal is to decrease negative affect when gaze is directed away from the emotionally evocative elements (Manera et al., 2014; van Reekum et al., 2007). In all likelihood, individuals monitor their current state, willfully search and redirect their attention to emotionally arousing elements of a complex image, and engage with a narrative to achieve the regulatory goal, reflected in MD. It is worth noting that whilst the neutral condition was the least cognitively demanding, the condition was still associated with a robust BOLD response, albeit at the lowest magnitude of all conditions. Most images included in the IAPS (Lang et al., 2008) are visually complex, even the more neutral images, requires visual scanning and decoding of meaning, likely reflected in MD. Indeed, in this study, the image sets selected were matched as much as possible on visual complexity and aspects such as the social nature of the image.

### Response (RESP)

The response mode is commonly observed in tasks requiring a motor response (Mascarenhas & Woodward, 2025). Our study is no exception, as participants were asked to press a button to provide a rating of the image when a rating screen appeared following image offset. Both the spatial map and the time course of the hemodynamic response function support the notion that the ratings engaged this brain network. The neutral image condition was associated with the highest and fastest hemodynamic response, likely reflecting the relative ease of evaluating neutral images as “neutral” compared to the negative images, given the gradation in rating options for “negative”. In its purest form, the RESP mode would not be load-dependent or otherwise show condition differences (Mascarenhas & Woodward, 2025). Also, with one-handed – in most cases right-handed - responses, the pattern would largely be left-lateralised. In our study, activation patterns were observed in both left and right hemisphere – albeit with higher intensity activation on the left – and demonstrated condition differences as summarised above. Furthermore, this component included activation in areas commonly associated with visual attention. Providing a rating involves making a judgment as to the subjective intensity of negative affect, and mapping the judgment onto visually displayed four response categories corresponding to 4 buttons on the response box. Thus, motor planning and responding are integrated with visual attention and decision-making. These processes combined likely drove the anatomical pattern obtained, and the variability in the temporal hemodynamic response pattern observed across the regulation conditions.

### Re-evaluation (RE-EV)

The classification of this component as “re-evaluation” out of 12 cognitive mode templates is particularly striking and adds to findings recently reported by Morawetz and colleagues (2020) and Bo and colleagues (2024). As highlighted by Redway and colleagues (2024), the re-evaluation mode template was derived from maps obtained during purely cognitive tasks: an adapted Stroop task requiring task switching, and two versions of a “bias against disconfirmatory evidence” task, a task in which participants decide whether partial information presented contradicted their expectation or belief. The mode template thus comprised spatial brain maps from tasks which were non-affective and different in nature from the reappraisal task used here, yet converged on the need to evaluate new information against a current mode. Indeed, to be effective, reappraisal commonly requires the override of a previous interpretation or change in belief (e.g. Troy et al., 2018), thereby supporting the presence of this mode for the reappraisal task requirement. The spatial map of the template is characterised by extensive involvement of medial and lateral frontal and parietal regions (BA 9/46/40), middle and inferior temporal regions (BA20/21), cerebellum and caudate. The component map from our study largely converged with the template map.

The temporal profiles of the regulation conditions indicated that the peak of RE-EV occurred at 15 s, which is after the peak observed for MD (11 s) and RESP (17 s). The HDRs also demonstrated the highest engagement of RE-EV for the “decrease” condition, followed by the “increase” then the “attend” condition. Our prior work, using deconvolution to estimate the task-evoked BOLD response (e.g. Urry et al., 2009; van Reekum et al., 2007), suggested lower engagement of several brain regions, reflected also in some autonomic measures of effort and arousal, for the decrease relative to the increase condition. Eye tracking data highlighted that lower brain activation for the decrease condition could be related to gaze aversion, or attentional disengagement, per our discussion above (under MD). The RE-EV findings suggest that the decrease condition involves most activity in the re-evaluation network, dispelling suspicion that participants mentally “switched off” during this condition. Instead, highest engagement of RE-EV for the decrease condition likely reflects incongruency between current state (a ‘bottom-up’ negative affect response to a stimulus appraised as aversive) and goal state (decreasing the negative affective response), requiring switching ongoing thoughts and feelings, not unlike integrating evidence that disconfirms a previous belief, c.f. tasks that formed the basis of the RE-EV mode (Redway, Arreaza, et al., 2025). The flat profile for the neutral condition underscores the distinct active characterisation of this cognitive mode, given that the task instruction was invariable for the neutral images (always “maintain attention to the image”).

### Default Mode B (DMB)

Any mental task is associated with fMRI patterns of relative de-activation in brain regions comprising the “default mode network” (Raichle et al., 2001). The reappraisal task is no exception. The template maps of fMRI-CPCA distinguish between two default modes, i.e. default mode A and B (Ni et al., 2025; Redway, Jian, et al., 2025). The typical spatial map and temporal profile of default mode B (DMB) corresponds to the oft-reported “default mode network” (Raichle et al., 2001) observed during self-reflective processes and “rest”. Default mode A (DMA) is more distinct and relatively novel. Whilst DMA frequently co-occurs as the deactivation component with, and typically comprises a mirror temporal profile to, the Language (LAN) mode, DMB is commonly reciprocally associated with MD. For this study, the spatial map of the component demonstrated overlap with both DMA and DMB, and fits DMB only moderately well (Fisher’s Z = 0.5). In addition, clusters in lateral OFC and left inferior frontal gyrus (pars triangularis) were part of the component here, but are not part of the DMA nor DMB template modes. Future research using fMRI-CPCA will help determine the extent to which these clusters are unique to tasks with an emotion element or shared with a different subset of tasks.

The BOLD changes demonstrated the most complex response pattern for the increase condition. Compared to the other conditions, the BOLD response for the increase condition started to rise later, and time to peak was delayed. Whilst in prior work, DMB reciprocally relates to MD, the temporal profiles of the conditions in this study suggests the patterns are more akin to RESP and, given the timing of the peaks around 15 s, RE-EV. This observation suggests that DMB was at least partly modulated by task-related factors beyond those already identified. Reappraisal involves a variety of processes to achieve the regulatory goal (of increasing or decreasing experienced emotion), and self-reflective processes, including semantic processes to give meaning to the images shown, are likely engaged when performing this task. Such processes could therefore have modulated the time courses of deactivation in DMB during reappraisal. Note that this interpretation of the finding is highly speculative at this stage. Future research with an emotion regulation-relevant task where such processes are better controlled could help to further disentangle the functional significance of default mode patterns observed, but our findings do highlight a possible flexibility of responses of DMB depending on task demands.

Using different analytic approaches, Morawetz and colleagues (2020) identified two meta-analytic groupings which matched cognitive task maps derived from working memory, reasoning, explicit memory and inhibition, and language, in the BrainMap database. Bo and colleagues (2024) separated a network that was selectively involved in reappraisal to regulate emotion from a network encompassing appraisal processes common in emotion elicitation and regulation. FMRI-CPCA decomposes the BOLD response both in space and time, providing spatial maps with temporal signatures that can further aid in interpreting the functional significance. Taking into account the temporal profiles and correlating the spatial component maps to previously identified templates of cognitive modes, the findings presented here suggest that “multiple demand” and “re-evaluation” modes are most relevant to reappraisal, more than “Language” or “Maintaining Internal Attention” modes which cover processes previously suggested to be relevant to reappraisal (e.g. Buhle et al., 2014). While such reverse inferences remain speculative until direct, causal evidence is obtained, our findings extend the results obtained by Morawetz and colleagues, and Bo and colleagues, highlighting that there are multiple, largely cortical, modes involved in reappraisal, which could be further probed in subsequent studies.

In conclusion, these findings highlight that the cortical networks most strongly engaged by reappraisal are linked to cognitive processes of “multiple demand” and “re-evaluation”. Furthermore, suggesting that increasing and decreasing negative affect are not two sides of the same coin, the regulatory goal was reflected in different response patterns in multiple demand and re-evaluation, as well as in deactivation of the default mode network. Employing a different analytic approach, this study furthers recent findings by Bo and colleagues regarding shared and distinct network contributions to reappraisal (emotion regulation) and appraisal (emotion generation) by 1) identifying key cognitive modes involved in reappraisal and 2) underlining the role of regulatory goal in the engagement of different brain networks. These novel findings help us to better understand the myriad of neurocognitive systems involved in emotion regulation, and identify targets for cognitive and neuroscience-based interventions to promote mental health and wellbeing.

## Data and Code Availability

The MRI data are publicly available on OpenNeuro:

https://openneuro.org/datasets/ds002620

Code for fMRI-CPCA is available at: https://www.nitrc.org/projects/fmricpca

## Author Contributions

(mandatory)

Carien van Reekum: Conceptualisation, Methodology, Supervision, Writing, Project Administration, Funding Acquisition. Emma Tupitsa: Methodology, Validation, Formal Analysis, Visualisation, Writing—Review & Editing, Project Administration. Will Lloyd: Methodology, Visualisation, Writing—Review & Editing, Project Administration. Ava Momeni: Formal Analysis, Visualisation, Writing – Review & Editing; John Shahki: Formal Analysis, Visualisation, Writing – Review & Editing; Eva Feredoes: Conceptualisation, Writing—Review & Editing, Funding Acquisition. Todd Woodward: Conceptualisation, Methodology, Software, Validation, Formal Analysis, Visualisation, Writing—Review & Editing

## Acknowledgements

Many thanks to Charlie Wan of the Woodward lab who helped with the component classification and HDR statistics, Karin Joanknecht who helped to collect the original data, and all our participants for their interest and for volunteering in our research. Will Lloyd is now at the University of Manchester; Eva Feredoes is now at Stanford University.

## Supplementary Material

(created during production as a web link to online material)

## References

American Psychiatric Association. (2022). Diagnostic and Statistical Manual of Mental Disorders (DSM-5-TR). American Psychiatric Association Publishing. 10.1176/appi.books.9780890425787

Beckmann, C. F., & Smith, S. M. (2004). Probabilistic Independent Component Analysis for Functional Magnetic Resonance Imaging. IEEE Transactions on Medical Imaging, 23(2), 137–152. 10.1109/TMI.2003.822821

Bo, K., Kraynak, T. E., Kwon, M., Sun, M., Gianaros, P. J., & Wager, T. D. (2024). A systems identification approach using Bayes factors to deconstruct the brain bases of emotion regulation. Nature Neuroscience, 27(5), 975–987. 10.1038/s41593-024-01605-7

Braunstein, L. M., Gross, J. J., & Ochsner, K. N. (2017). Explicit and implicit emotion regulation: A multi-level framework. Social Cognitive and Affective Neuroscience, 12(10), 1545–1557. 10.1093/scan/nsx096

Buhle, J. T., Silvers, J. A., Wager, T. D., Lopez, R., Onyemekwu, C., Kober, H., Weber, J., & Ochsner, K. N. (2014). Cognitive Reappraisal of Emotion: A Meta-Analysis of Human Neuroimaging Studies. Cerebral Cortex, 24(11), 2981–2990. 10.1093/cercor/bht154

Cox, R. W. (1996). AFNI: Software for analysis and visualization of functional magnetic resonance neuroimages. Comput. Biomed. Res., 29(3), 162–173.

De Voogd, L. D., & Hermans, E. J. (2022). Meta-analytic evidence for downregulation of the amygdala during working memory maintenance. Human Brain Mapping, 43(9), 2951–2971. 10.1002/hbm.25828

Duncan, J. (2010). The multiple-demand (MD) system of the primate brain: Mental programs for intelligent behaviour. Trends in Cognitive Sciences, 14(4), 172–179. 10.1016/j.tics.2010.01.004

Evora, M., & Woodward, T. S. (2025). Cognitive Mode Detectable with Task-Based fMRI: Initiation (INIT). In Percival, Chantal M. & T. S. Woodward (Eds), Cognitive Modes Detectable with Task-Based fMRI (pp. 207–229). PsyArXiv. 10.31234/osf.io/8mbqe_v3

Griffanti, L., Douaud, G., Bijsterbosch, J., Evangelisti, S., Alfaro-Almagro, F., Glasser, M. F., Duff, E. P., Fitzgibbon, S., Westphal, R., Carone, D., Beckmann, C. F., & Smith, S. M. (2017). Hand classification of fMRI ICA noise components. NeuroImage, 154, 188–205. 10.1016/j.neuroimage.2016.12.036

Gross, J. J. (1998). Antecedent- and response-focused emotion regulation: Divergent consequences for experience, expression, and physiology. Journal of Personality and Social Psychology, 74(1), 224–237. 10.1037/0022-3514.74.1.224

He, Z., Li, S., Mo, L., Zheng, Z., Li, Y., Li, H., & Zhang, D. (2023). The VLPFC-Engaged Voluntary Emotion Regulation: Combined TMS-fMRI Evidence for the Neural Circuit of Cognitive Reappraisal. The Journal of Neuroscience, 43(34), 6046–6060. 10.1523/JNEUROSCI.1337-22.2023

Jenkinson, M., Bannister, P., Brady, M., & Smith, S. (2002). Improved optimization for the robust and accurate linear registration and motion correction of brain images. NeuroImage, 17(2), 825–841.

Jenkinson, M., Beckmann, C. F., Behrens, T. E. J., Woolrich, M. W., & Smith, S. M. (2012). FSL. NeuroImage, 62(2), 782–790. 10.1016/j.neuroimage.2011.09.015

Kohn, N., Eickhoff, S. B., Scheller, M., Laird, A. R., Fox, P. T., & Habel, U. (2014). Neural network of cognitive emotion regulation—An ALE meta-analysis and MACM analysis. NeuroImage, 87, 345–355. 10.1016/j.neuroimage.2013.11.001

Lang, P., Bradley, M., & Cuthbert, B. (2008). International affective picture system (IAPS): Affective ratings of pictures and instruction manual.

Lloyd, W. K., Morriss, J., Macdonald, B., Joanknecht, K., Nihouarn, J., & Van Reekum, C. M. (2021). Longitudinal change in executive function is associated with impaired top-down frontolimbic regulation during reappraisal in older adults. NeuroImage, 225, 117488. 10.1016/j.neuroimage.2020.117488

Manera, V., Samson, A. C., Pehrs, C., Lee, I. A., & Gross, J. J. (2014). The eyes have it: The role of attention in cognitive reappraisal of social stimuli. Emotion, 14(5), 833–839. 10.1037/a0037350

Mascarenhas, M., & Woodward, T. S. (2025). Cognitive Mode Detectable with Task-Based fMRI: Response (RESP). In Percival, Chantal M. & T. S. Woodward (Eds), Cognitive Modes Detectable with Task-Based fMRI (pp. 310–337). PsyArXiv. 10.31234/osf.io/8mbqe_v3

Metzak, P., Feredoes, E., Takane, Y., Wang, L., Weinstein, S., Cairo, T., Ngan, E. T. C., & Woodward, T. S. (2011). Constrained principal component analysis reveals functionally connected load-dependent networks involved in multiple stages of working memory. Human Brain Mapping, 32(6), 856–871. 10.1002/hbm.21072

Momeni, A., Addis, D. R., Feredoes, E., Klepel, F., Rasheed, M. M., Chinchani, A. M., Koussis, N. C., & Woodward, T. S. (2025). Functional Brain Networks Underlying Autobiographical Event Simulation: An Update. Journal of Cognitive Neuroscience, 37(6), 1083–1146. 10.1162/jocn_a_02305

Morawetz, C., Bode, S., Derntl, B., & Heekeren, H. R. (2017). The effect of strategies, goals and stimulus material on the neural mechanisms of emotion regulation: A meta-analysis of fMRI studies. Neuroscience & Biobehavioral Reviews, 72, 111–128. 10.1016/j.neubiorev.2016.11.014

Morawetz, C., Riedel, M. C., Salo, T., Berboth, S., Eickhoff, S. B., Laird, A. R., & Kohn, N. (2020). Multiple large-scale neural networks underlying emotion regulation. Neuroscience & Biobehavioral Reviews, 116, 382–395. 10.1016/j.neubiorev.2020.07.001

Ni, Y. Q. Y., Redway, S., Momeni, A., Momeni, A., Jian, L., Yip, L., & Woodward, T. S. (2025). Cognitive Mode Detectable with Task-Based fMRI: Default Mode B (DMB). In C. M. Percival & T. S. Woodward (Eds), Cognitive Modes Detectable with Task-Based fMRI (pp. 360–395). 10.31234/osf.io/8mbqe_v3

Ochsner, K. N., Ray, R. D., Cooper, J. C., Robertson, E. R., Chopra, S., Gabrieli, J. D., & Gross, J. J. (2004). For better or for worse: Neural systems supporting the cognitive down- and up-regulation of negative emotion. NeuroImage, 23(2), 483–499.

Ochsner, K. N., Silvers, J. A., & Buhle, J. T. (2012). Functional imaging studies of emotion regulation: A synthetic review and evolving model of the cognitive control of emotion. Annals of the New York Academy of Sciences, 1251(1). 10.1111/j.1749-6632.2012.06751.x

Percival, C. M., & Woodward, T. S. (2025). Cognitive Modes Detectable with Task-Based fMRI. PsyArXiv. 10.31234/osf.io/8mbqe_v3

Percival, C. M., Zahid, H. B., & Woodward, T. S. (2020). CNoS-Lab/Woodward_Atlas (Version 1.0.3) [Computer software]. Zenodo. 10.5281/ZENODO.4281551

Raichle, M. E., MacLeod, A. M., Snyder, A. Z., Powers, W. J., Gusnard, D. A., & Shulman, G. L. (2001). A default mode of brain function. Proceedings of the National Academy of Sciences, 98(2), 676–682. 10.1073/pnas.98.2.676

Redway, S., Arreaza, L., Shahki, J., Zeng, E., Tsang, J., & Woodward, T. S. (2025). Cognitive Mode Detectable with Task-Based fMRI: Re-Evaluation (RE-EV). In Percival, Chantal M. & T. S. Woodward (Eds), Cognitive Modes Detectable with Task-Based fMRI (pp. 152–180). PsyArXiv. 10.31234/osf.io/8mbqe_v3

Redway, S., Jian, L., Yip, L., Tsang, J., & Woodward, T. S. (2025). Cognitive Mode Detectable with Task-Based fMRI: Default Mode A (DMA). In Percival, Chantal M. & T. S. Woodward (Eds), Cognitive Modes Detectable with Task-Based fMRI (pp. 338–359). PsyArXiv. 10.31234/osf.io/8mbqe_v3

Roes, M. M., Yin, J., Taylor, L., Metzak, P. D., Lavigne, K. M., Chinchani, A., Tipper, C. M., & Woodward, T. S. (2020). Hallucination-Specific structure-function associations in schizophrenia. Psychiatry Research: Neuroimaging, 305, 111171. 10.1016/j.pscychresns.2020.111171

Rorden, C., & Brett, M. (2000). Stereotaxic Display of Brain Lesions. Behavioural Neurology, 12(4), 191–200. 10.1155/2000/421719

Sanford, N., Whitman, J. C., & Woodward, T. S. (2020). Task-merging for finer separation of functional brain networks in working memory. Cortex, 125, 246–271. 10.1016/j.cortex.2019.12.014

Shashidhara, S., Assem, M., Glasser, M. F., & Duncan, J. (2024). Task and stimulus coding in the multiple-demand network. Cerebral Cortex, 34(7), bhae278. 10.1093/cercor/bhae278

Smith, S. M. (2002). Fast robust automated brain extraction. Human Brain Mapping, 17(3), 143–155. 10.1002/hbm.10062

Smith, S. M., Jenkinson, M., Woolrich, M. W., Beckmann, C. F., Behrens, T. E. J., Johansen-Berg, H., Bannister, P. R., De Luca, M., Drobnjak, I., Flitney, D. E., Niazy, R. K., Saunders, J., Vickers, J., Zhang, Y., De Stefano, N., Brady, J. M., & Matthews, P. M. (2004). Advances in functional and structural MR image analysis and implementation as FSL. NeuroImage, 23, Supplement 1, S208– S219. 10.1016/j.neuroimage.2004.07.051

Thomas Yeo, B. T., Krienen, F. M., Sepulcre, J., Sabuncu, M. R., Lashkari, D., Hollinshead, M., Roffman, J. L., Smoller, J. W., Zöllei, L., Polimeni, J. R., Fischl, B., Liu, H., & Buckner, R. L. (2011). The organization of the human cerebral cortex estimated by intrinsic functional connectivity. Journal of Neurophysiology, 106(3), 1125–1165. 10.1152/jn.00338.2011

Troy, A. S., Shallcross, A. J., Brunner, A., Friedman, R., & Jones, M. C. (2018). Cognitive reappraisal and acceptance: Effects on emotion, physiology, and perceived cognitive costs. Emotion, 18(1), 58–74. 10.1037/emo0000371

Tupitsa, E., Egbuniwe, I., Lloyd, W. K., Puertollano, M., Macdonald, B., Joanknecht, K., Sakaki, M., & Van Reekum, C. M. (2023). Heart rate variability covaries with amygdala functional connectivity during voluntary emotion regulation. NeuroImage, 274, 120136. 10.1016/j.neuroimage.2023.120136

Tzourio-Mazoyer, N., Landeau, B., Papathanassiou, D., Crivello, F., Etard, O., Delcroix, N., Mazoyer, B., & Joliot, M. (2002). Automated Anatomical Labeling of Activations in SPM Using a Macroscopic Anatomical Parcellation of the MNI MRI Single-Subject Brain. NeuroImage, 15(1), 273–289. 10.1006/nimg.2001.0978

Urry, H. L., van Reekum, C. M., Johnstone, T., & Davidson, R. J. (2009). Individual differences in some (but not all) medial prefrontal regions reflect cognitive demand while regulating unpleasant emotion. NeuroImage, 47(3), 852–863. 10.1016/j.neuroimage.2009.05.069

Urry, H. L., van Reekum, C. M., Johnstone, T., Kalin, N. H., Thurow, M. E., Schaefer, H. S., Jackson, C. A., Frye, C. J., Greischar, L. L., Alexander, A. L., & others. (2006). Amygdala and ventromedial prefrontal cortex are inversely coupled during regulation of negative affect and predict the diurnal pattern of cortisol secretion among older adults. Journal of Neuroscience, 26(16), 4415.

van Reekum, C. M., Johnstone, T., Urry, H. L., Thurow, M. E., Schaefer, H. S., Alexander, A. L., & Davidson, R. J. (2007). Gaze fixations predict brain activation during the voluntary regulation of picture-induced negative affect. Neuroimage, 36(3), 1041–1055.

Wadden, K. P., Woodward, T. S., Metzak, P. D., Lavigne, K. M., Lakhani, B., Auriat, A. M., & Boyd, L. A. (2015). Compensatory motor network connectivity is associated with motor sequence learning after subcortical stroke. Behavioural Brain Research, 286, 136–145. 10.1016/j.bbr.2015.02.054

Wager, T. D., Davidson, M. L., Hughes, B. L., Lindquist, M. A., & Ochsner, K. N. (2008). Prefrontal-Subcortical Pathways Mediating Successful Emotion Regulation. Neuron, 59(6), 1037–1050. 10.1016/j.neuron.2008.09.006

Wang, Z., Shahki, J., Jian, L., & Woodward, T. S. (2025). Cognitive Mode Detectable with Task-Based fMRI: Multiple Demand (MD). In C. M. Percival & T. S. Woodward (Eds), Cognitive Modes Detectable with Task-Based fMRI (pp. 181–206). PsyArXiv. https://osf.io/preprints/psyarxiv/8mbqe_v3

Wong, S. T. S., Goghari, V. M., Sanford, N., Lim, R., Clark, C., Metzak, P. D., Rossell, S. L., Menon, M., & Woodward, T. S. (2020). Functional brain networks involved in lexical decision. Brain and Cognition, 138, 103631. 10.1016/j.bandc.2019.103631

Woodward, T. S., Cairo, T. A., Ruff, C. C., Takane, Y., Hunter, M. A., & Ngan, E. T. C. (2006). Functional connectivity reveals load dependent neural systems underlying encoding and maintenance in verbal working memory. Neuroscience, 139(1), 317–325. 10.1016/j.neuroscience.2005.05.043

Yarkoni, T., Poldrack, R. A., Nichols, T. E., Van Essen, D. C., & Wager, T. D. (2011). Large-scale automated synthesis of human functional neuroimaging data. Nature Methods, 8(8), 665–670. 10.1038/nmeth.1635

